# Modelling the tonotopic map using a two-dimensional array of neural oscillators

**DOI:** 10.1101/2022.02.21.481388

**Authors:** Dipayan Biswas, V. Srinivasa Chakravarthy, Asit Tarsode

## Abstract

We present a model of a tonotopic map known as the Oscillatory Tonotopic Self-Organizing Map (OTSOM). It is a 2-dimensional, self-organizing array of Hopf oscillators, capable of performing a Fourier-like decomposition of the input signal. While the rows in the map encode the input phase, the columns encode frequency. Although Hopf oscillators exhibit resonance to a sinusoidal signal when there is a frequency match, there is no obvious way to also achieve phase tuning. We propose a simple method by which a pair of Hopf oscillators, unilaterally coupled through a coupling scheme termed as modified power coupling, can exhibit tuning to the phase offset of sinusoidal forcing input. The training of OTSOM is performed in 2 stages: while the frequency tuning is adapted in stage 1, phase tuning is adapted in stage 2. Earlier tonotopic map models have modeled frequency as an abstract parameter unconnected to any oscillation. By contrast, in OTSOM, frequency tuning emerges as a natural outcome of an underlying resonant process. The OTSOM model can be regarded as an approximation of the tonotopic map found in the primary auditory cortices of mammals, particularly exemplified in the studies of echolocating bats.

## 1 Introduction

The discovery of cortical brain maps in mammalian brains is perhaps one of the first milestones in our understanding of how the brain generates representations of the world. Visual research had discovered a rich hierarchy of maps of various sub-modalities of vision (orientation, curvature, color, and even complex objects) in various visual cortical areas extended over the occipital, parietal, and temporal lobes (Hadjikhani et al., 1998; Hubel & Wiesel, 1959; Wandell et al., 2007; Yue et al., 2020). A similar network of maps of somatotopy was found in the somatosensory areas of the postcentral gyrus, and posterior parietal cortex (Penfield, 1937). These studies have placed on a firm foundation the understanding that sensory information in the brain is often laid out in the form of a system of topographic maps. However, efforts to establish a similar map structure underlying auditory processing - popularly referred as tonotopic maps, - are met with considerable challenges.

Tonotopy begins in the inner ear, in the hair cells laid out along the length of the basilar membrane inside the cochlea (Bekesy, 1949). Parts of the basilar membrane respond to different frequencies, with the tuning frequency increasing in the apex to the base direction (Ruggero, 1992). Thus, there is a well-established tonotopy in the cochlea, sometimes also referred to as cochleotopy. Beyond the cochlea, there is a hierarchy of areas along the auditory pathway (Clopton et al., 1974; Ehret & Romand, 1996; Palmer & Rees, 2010). Although there is a general agreement that what is mapped in tonotopic maps is the frequencies, other auditory parameters like sound intensity, tuning bandwidth are also explored (Boynton et al., 2015; Schreiner & Sutter, 1992).

Earliest studies on tonotopy focused on frequency tuning, treating it as one of the primary features if not the sole defining feature of auditory response. Merzenich et al., 2018 found a systematic representation of cochlea within the primary auditory cortex of cats. It was observed that frequency bands of the input stimuli are mapped onto rectilinear strips in the auditory cortex. Similar observations were made in the auditory cortex of the grey squirrel (Michael M. Merzenich et al., 1976). Investigations of the auditory cortex in owl monkeys have discovered a central area with orderly mapping of audible frequencies, circumscribed by areas where neurons show more complex responses than frequency tuning (Imig & Adrian, 1977). However, the exact nature of the complex responses was not elaborated in the last study.

A hierarchically organized network of areas with complex information processing properties was discovered subsequently in the auditory cortex of the bat (N. Suga, 1990). Contrary to popular belief, bats are not visually blind, though the extent of visual capacity varies with different subspecies of bats. But bats predominantly depend on echolocation to navigate through the spatial world. Bats emit ultrasound pulses in the frequency range of tens of kilo Hertz, and interpret the spatial world from the echoes returned by the environment. Whereas the delay between the emitted and the received pulse reveals distance, doppler shift in the echo reveals the relative velocity between the echolocating bat and a target. Pioneering studies of the bat’s auditory system by Alvin Novick, James Simmons, Nobuo Suga, and others had revealed that these complex auditory functions of the bat are subserved by a well-developed auditory system (Bates et al., 2011; Novick & Vaisnys, 1964; Simmons, 2012; N. Suga, 1990; Nobuo Suga et al., 1997).

Studies by Nobuo Suga and colleagues with the mustached bat had described an elaborate network of auditory cortical areas with an intrinsic hierarchy not very different from that of the primate visual system (Hubel & Wiesel, 1959). The mustached bat emits composite pulses that have an initial Constant Frequency (CF) section terminated by a Frequency Modulated (FM) section. There is a cortical region in which neurons respond only to certain combinations of frequencies and amplitudes of echoes. There is a region where neurons respond only to frequency differences between the emitted pulse and its echoes, probably encoding Doppler shift. In another region there are neurons that respond to the time delay between the emitted pulse and the echo, perhaps encoding the distance to the target. The gains obtained from the study of the bat’s auditory system are not yet fully exploited in unravelling the auditory architecture of the brains of higher mammals and humans.

In the domain of computational modelling, one of the earliest tonotopic map models used a Self-Organizing Map (SOM) model to model the auditory cortex of mustached bat (Ritter et al., 1992). The model adopted a simplistic view of the organization of the bat’s auditory cortex – that the input frequencies are mapped along a rectilinear strip of the cortex – and shows how such a mapping can be realized using a rectangular SOM model. A key limitation of the model is the representation of frequency as an explicit scalar variable and not as an implicit temporal property of an ongoing oscillation. The SOM approach to modelling tonotopy was extended to construct a model of a “phonetic typewriter” (Kohonen, 1988). Palakal et al., 1995 presented a tonotopic map model also based on the SOM approach, describing neural tuning to both frequency and delay. Here too frequency and delay are explicitly represented as scalar variables, and not as implicit temporal properties of a signal.

Models tend to make simplifying assumptions of the processes they aim to model. But it is rather unnatural to model frequency as simply a number without explicitly modelling the oscillation that the frequency refers to. A tonotopic map is primarily a response to tones which are oscillations. Oscillatory activities are found at all levels in the auditory pathway, from cochlea to inferior colliculus to higher auditory cortical areas. There is a considerable body of literature that examines the application of nonlinear oscillators to describe responses of the auditory system at various levels to periodic stimulus(Frank Julicher et al., 2001; Fredrickson-hemsing et al., 2012; Kim & Large, 2015; Laudanski et al., 2010; Meddis & Lowel P. O’Mard, 2006; Víctor M. Eguíluz & Ospeck, 2000). However, these are single-unit models of oscillation and not map models.

From the aforementioned quick review of auditory response models, we understand that there are tonotopic map models that do not explicitly model the underlying oscillation, and there are oscillatory models at single unit level that are not extended to map models. Thus, the challenge of constructing a tonotopic map model of nonlinear oscillators is still unrealized, which becomes the motivation of the present work.

We present a tonotopic map model consisting of a 2-dimensional array of nonlinear oscillators. Specifically, we choose the Hopf oscillators since these oscillators have been extensively used to model auditory responses (Farokhniaee et al., 2020; Frank Julicher et al., 2001; Fredrickson-hemsing et al., 2012; Kim & Large, 2015; Large et al., 2010). In the following methods section, we have presented the dynamics of the OTSOM model along with the dynamical analysis of a single unit of the OTSOM model and the modified power coupling strategy along with the modified Hebbian learning rule to train it. The dynamical analysis of the two stages to train the characterizing frequencies and the phases of the model is presented thereafter. The numerical analysis from the unit level to the network level is presented in the subsequent results section.

## 2 Methods

The conventional self-organizing maps are known for their special characteristic of organizing internally represented features on a spatial scale. i.e., the neurons representing similar abstract features in the input data organize themselves spatially close to each other through competitive learning (Kohonen, 1998). Typically, SOM models perform dimensionality reduction of a high dimensional input vector by projecting it onto a low dimensional spatial map space. In other words, the input data points located nearby in the *N*-dimensional Euclidean space get mapped onto nearby neurons in the map. The map space can be maximum up to 3 dimensional as it is not easily visualized beyond 3 dimensions. Also, brain maps are typically 2-dimensional, referring typically to cortical sheets of neurons, or mildly 3-dimensional, if the small cortical thickness is included.

The objective of our modelling study is to propose a dynamical self-organizing map model which can organize the features of complex sinusoidal signals on a 2-dimensional grid of nonlinear oscillators. Signals of any duration have a static representation in Fourier space or frequency domain. The proposed model is capable of organizing features of complex sinusoidal signals such as frequency and phase offset, in terms of the parameters of intrinsic dynamics of single neural oscillator and the connectivity parameters during the training phase. During testing, the trained map model can represent the features of any composite signal with multiple frequency components.

### Oscillatory Tonotopic Self Organizing Map (OTSOM)

The Oscillatory Tonotopic Self Organizing Map (OTSOM) model consists of a 2D array of Hopf oscillators (Strogatz, 1994) - the “Cortical Array of Oscillators” (CAO) - operating at supercritical Hopf regime as described in fig. 1. A single isolated oscillator located apart from the CAO is interpreted as a subcortical oscillator, and labeled as Subcortical Reference Oscillator (SRO) since it serves as a reference for the phases of the cortical oscillators. Although there are no lateral interactions between the oscillators in CAO, the SRO projects unilateral, trainable connections to all the oscillators in CAO. Thus, each oscillator in CAO receives two inputs: 1) from the SRO (termed as *I_r_*) and, 2) from the external input (termed as *I_e_*) (fig. 1). The connections from the external input to the CAO oscillators have uniform fixed weights.

**Figure 1:**
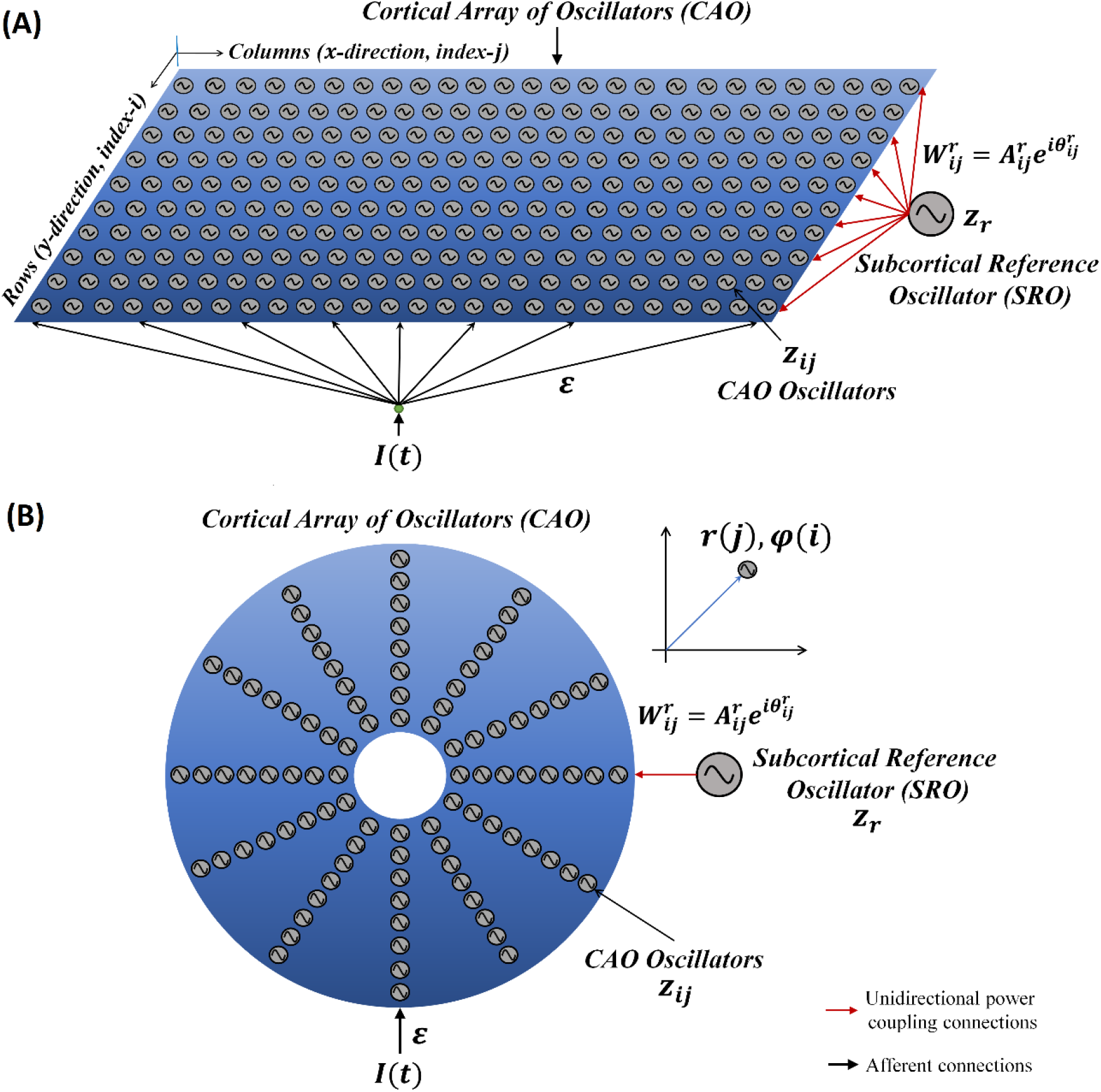
The network architecture of Oscillatory Tonotopic Self Organizing Map (OTSOM). The model principally contains two types of oscillators: an array of cortical oscillators (CAO) and a Subcortical Reference Oscillator (SRO). The SRO projects unilateral, power-coupling connections to the CAO oscillators. The external input perturbs the oscillators in CAO through uniform afferent connections. The CAO oscillators can be visualized to be organized either in a 2D array **(A)** or in concentric circular array **(B).** The radial and the angular position of a CAO oscillator in the concentric circular array is same as its position along x and y axis respectively in the 2D array organization of the cortical array.

Before going into more details of the dynamics of the OTSOM model let us first understand the dynamics of the constituting components of the OTSOM model. As mentioned before, the CAO oscillators and the SRO are Hopf oscillators. A single Hopf oscillator can be represented in Cartesian (eqns. 1a, b) and polar coordinates (eqns. 2a, b) respectively as follows:

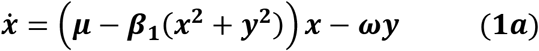

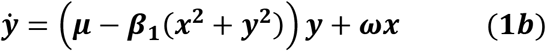

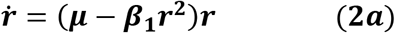

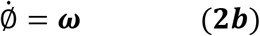

where, (*x,y*) are cartesian coordinate variables and (*r*, Ø) are polar coordinate variables; *ω* defines the angular velocity or the natural frequency of the oscillator. The parameters *μ* and *β*_1_ determine the dynamic regime of the Hopf oscillator: for *μ* = 0, *β*_1_ > 0 it operates in critical Hopf regime; for *μ* > 0, *β*_1_ > 0 it operates in supercritical Hopf regime and when *μ* = 0, *β*_1_ = 0 it is a simple harmonic oscillator (Kim & Large, 2015).

Combining x and y of eqn. (1) into a complex number, *z* = *x* + *iy*, Hopf oscillator dynamics can simply be represented on complex plane elegantly as:

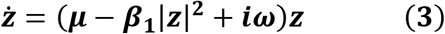

where, 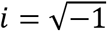. A pair of coupled Hopf oscillators can principally exhibit two types of dynamical phenomena: synchronization and entrainment.

A generalized definition of synchronization has been introduced in our previous study (Biswas et al., 2021), which states that any two oscillators can be claimed to be synchronized irrespective of their intrinsic oscillation frequencies if they maintain any of the following phase relationships constant, 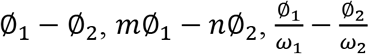; *m* and *n* are natural numbers. Whereas entrainment is the dynamical characteristic of an oscillator whilst the frequency of oscillation of the oscillator gradually changes from its natural frequency of oscillation to a new value when the oscillator is either coupled with another oscillator or perturbed by an external oscillatory input of a different frequency. Real valued symmetric coupling yields in phase (0°) oscillation for positive coupling, and out of phase (180°) oscillation for negative coupling, between two isochronous oscillators, whereas the same pair can phase-lock at any arbitrary phase difference if coupled through ‘complex coupling’ strategy (Biswas et al, 2021).

To produce phase-locked dynamics from a pair of oscillators with unequal natural frequencies requires a special kind of complex coupling strategy labelled as ‘power coupling’ (Biswas et al., 2021). A pair of oscillators coupled through power coupling is defined as:

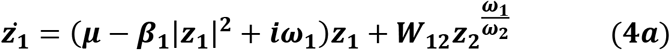

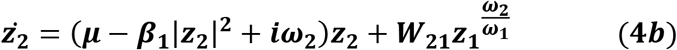

where, 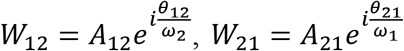 represent the complex power coupling coefficients on the feedforward and feedback branches. Considering *θ*_12_ = –*θ*_21_, and *A*_12_ = *A*_21_, it has been shown that the pair of oscillators can phase-lock at any of the solutions of the equation: 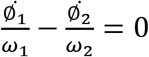 depending on the initial condition. The analytic solution of this problem gets increasingly complicated as the number of oscillators in the network increases. Another limitation of the original power-coupling strategy is that it does not ensure synchronization while the coupled oscillators are entrained to a new frequency of oscillation.

Considering a simplified scenario of a pair of Hopf oscillators coupled through weak unilateral power coupling, the one receiving input from the other oscillator through power coupling is being perturbed by a strong complex sinusoidal external input signal with frequency close to the natural frequency of the oscillator as depicted in fig. 2A.

**Figure 2:**
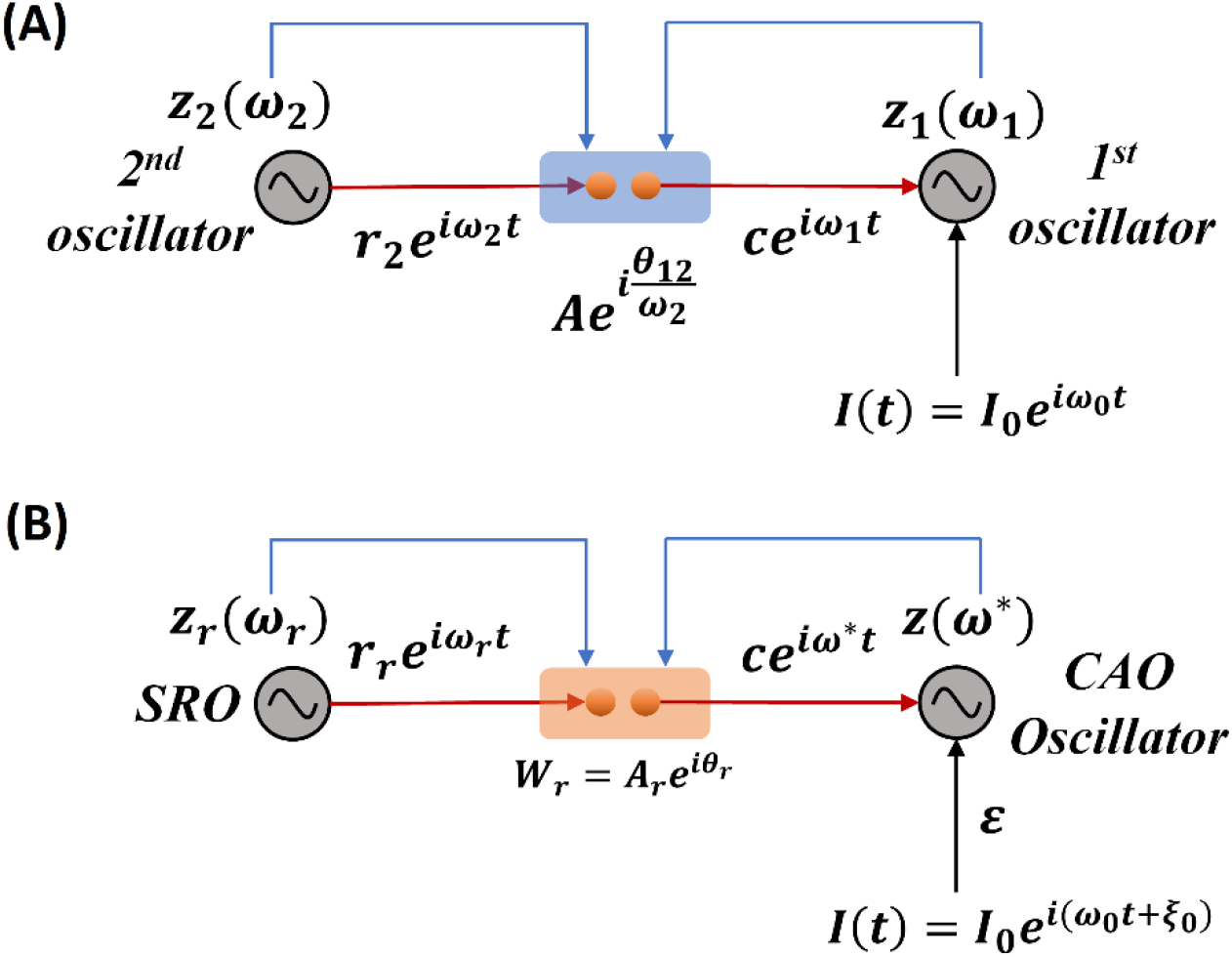
**(A)** The schematic diagram of a pair of Hopf oscillators, unilaterally coupled through conventional power coupling strategy. Here *c* is typically 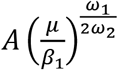 at steady state. **(B)** The fundamental building block of the OTSOM model. The single CAO oscillator receives inputs from two sources: one from the SRO and the other from the external input signal. The two differences between these two frameworks are the coupling coefficients; the natural frequency of the SRO is eliminated from the denominator of the angle of the coupling coefficient, and the second being the actual frequency of oscillation of the CAO oscillator is used to readjust the complex activation of the SRO (subplot A) instead of the natural frequency of the post-synaptic oscillator (subplot B).

We will now try to show the pair of oscillators with an external sinusoidal input as shown in fig. 2A, poses a new difficulty in phase-locking not faced by a pair of oscillators with power coupling (eqns. 4a,b). Consider the situation in fig. 2A, where the 2^nd^ Oscillator sends a unilateral power coupling connection to the 1^st^ Oscillator, which in addition receives a complex sinusoidal signal of frequency *ω*_0_ as external input. Note that though the output of the 2^nd^ Oscillator has the frequency *ω*_2_, after the power coupling connection, the signal frequency changes to *ω*_1_. Thus the 1^st^ Oscillator receives two sinusoidal inputs – of frequencies *ω*_0_ and *ω*_1_. Therefore, the 1^st^ Oscillator does not simply entrain to the external input, due to the interference from the 2^nd^ Oscillator. In order to fix this problem, it turns out that we need a more general power coupling rule than the original one.

### Modified Power Coupling

To make the original power-coupling rule (Biswas et al 2021) more generalized, and to ensure synchronization in a pair of power-coupled oscillators even after entrainment, a modified version of the power-coupling strategy is proposed. For a pair of bilaterally coupled Hopf oscillators, the modified power-coupling mechanism is given as,

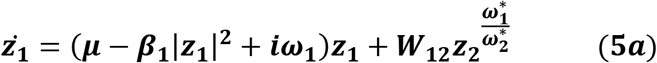

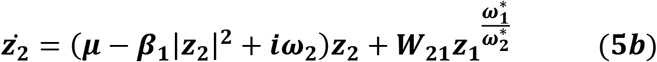

where, 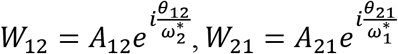, the only modification being 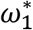 and 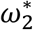 are actual frequencies of oscillation instead of natural frequencies. A new set of dynamic equations (eqns. 8a,b) define 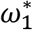 and 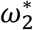. The overall dynamics of a pair of Hopf oscillators coupled through modified power-coupling is represented in polar coordinates as follows.

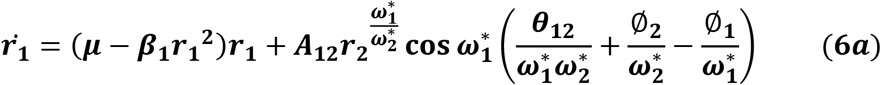

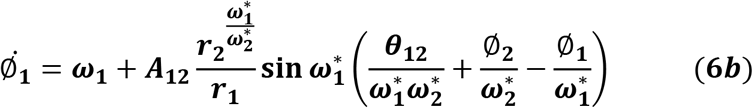

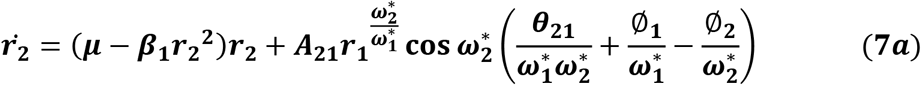

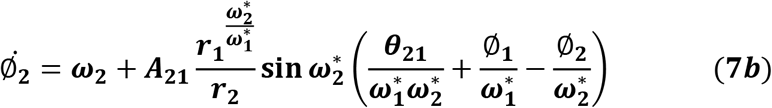

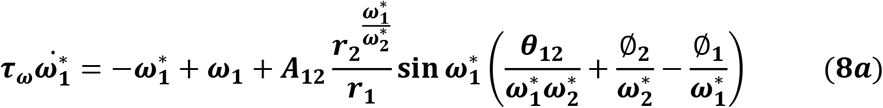

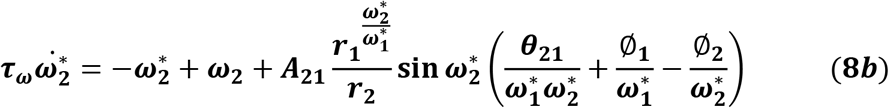

where, *τ_ω_* is the time constant for 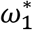 and 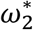. It is evident that without eqns. 8a and 8b, the modified power-coupling is functionally the same as the conventional one as *ω*\* remains *ω*. It can also be observed that neither of the oscillators can get entrained to a new frequency of oscillation, which makes a pair of bilaterally coupled Hopf oscillators through modified power coupling functionally the same as a pair of bilaterally coupled Hopf oscillators through conventional power coupling (refer to Appendix 1).

In the subsequent sections, the details of the intrinsic dynamics and the training framework of OSTOM model is described. The dynamics of a typical CAO oscillator is given as,

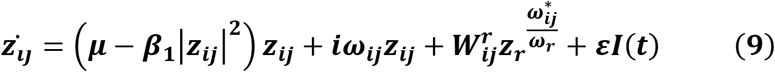

The first two terms on the RHS denote the intrinsic dynamics of the Hopf oscillator, the third term represents the input from SRO 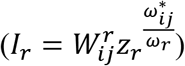, the fourth term represents the aggregate external input (*I_e_* = *εI*(*t*)), where *I*(*t*) is the actual external input. Note that only the SRO input is given via modified power coupling whereas the external input *I*(*t*) is presented directly with a multiplicative factor, ε.

Similarly, the dynamics of the SRO is given as,

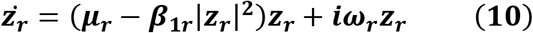

The activation of the CAO oscillator at location (*i,j*) is defined as, 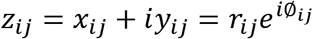; similarly, the activation of the SRO oscillator is 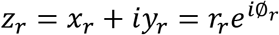. Intrinsic dynamics of the CAO oscillator is defined by the parameters *μ*, *β*_1_ and *ω_ij_*. Note that *μ* and *β*_1_ are the same for all CAO oscillators but *ω_ij_* is different. The intrinsic dynamics of the SRO oscillator is defined by *μ_r_, β*_1*r*_ and *ω_r_* parameters. 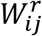 is the complex power-coupling weight from the reference oscillator to the oscillator at (*i,j*) in CAO, where 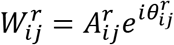.

The reason behind dropping the actual frequency of the presynaptic oscillator from the denominator of the angle of the complex coupling coefficient will be justified in the following sections.

The Cartesian and the polar coordinate representations are respectively.

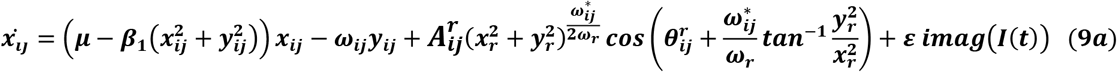

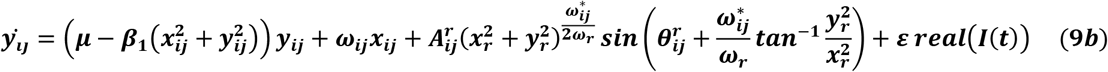

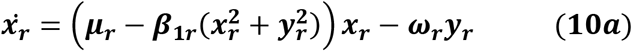

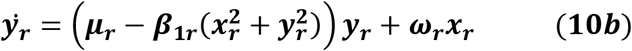

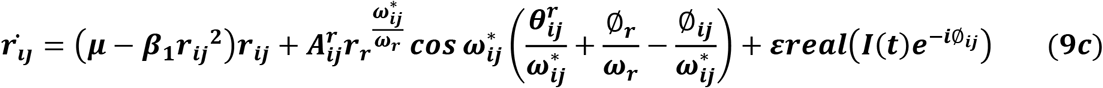

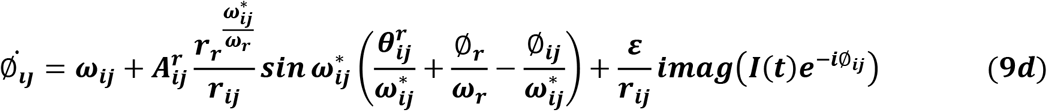

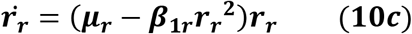

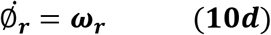

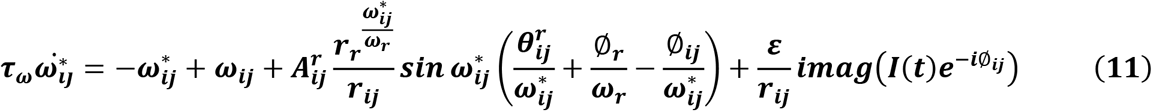

The uniform, real-valued, afferent connections from the external input (*I*(*t*)) to the CAO oscillators is *ε*. 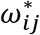 and 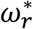 are the actual frequencies of the CAO oscillators and the SRO respectively. As 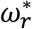 remains *ω_r_*, it is replaced with *ω*_r_. The actual frequency of oscillation of the CAO oscillator can be entrained to the frequency of the external perturbation. This entrainment property of the Hopf oscillator is utilized to realize the framework of the proposed model which will be discussed in detail in the later sections. Before elaborating the details of the training framework of the OTSOM model we are going the briefly analyze the intrinsic dynamical properties of the single unit (the SRO unilaterally coupled to a CAO oscillator) of the model.

#### Dynamical response of a single unit

The single unit, which is the fundamental building block of the whole OSTOM model is constituted of a single oscillator in the CAO which receives two inputs: 1) from the SRO via modified power coupling and, 2) from the external input. The SRO is coupled to a CAO oscillator unilaterally through a modified power coupling connection (eqns. 9,10,11). The output of the SRO, after passing through modified power-coupling connection, takes on the same frequency as that of the CAO oscillator it projects to. The CAO oscillator is also driven by external input signal through a real-valued afferent connection (fig. 2B). The external input is a complex sinusoidal signal.

We now investigate the steady state response of the single unit by analyzing (eqns. 12, 13, 14). Let the external input signal, 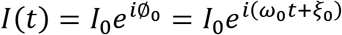 where *ω*_0_ and *ξ*_0_ are respectively the frequency and the phase offset of complex sinusoidal signal. Introducing the following variables,

- the relative phase of the CAO oscillator w.r.t the input, 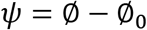,
- relative frequency of the CAO oscillator w.r.t the input, *Ω* = *ω* – *ω*_0_, and
- the normalized phase difference between the CAO oscillator and the SRO, 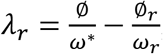, the eqns. 9,10,11 can be simplified to:

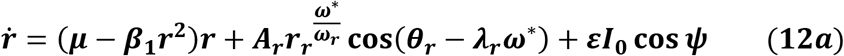

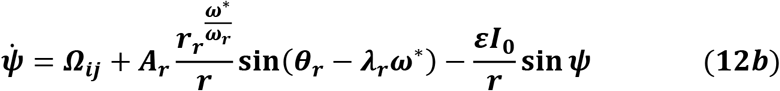

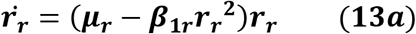

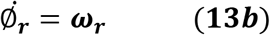

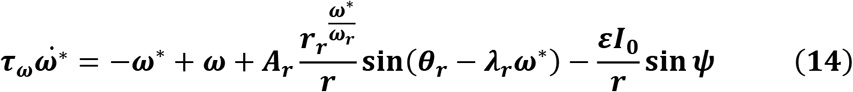

Under the special condition, 0 < *ε* ≪ 1, the single unit is equivalent to a pair of unidirectionally coupled oscillators through modified power coupling. There are two differences between the modified power coupling used under unilateral coupling scenario in the present study and the conventional power coupling proposed in (Biswas et al., 2021). In the conventional power coupling scheme of (Biswas et al 2021), the complex state of the oscillator is raised to the ratio of the *intrinsic* frequencies of the presynaptic and postsynaptic oscillators. In the modified power coupling proposed now, the exponent is the ratio between the *actual* frequency of oscillation of the post-synaptic oscillator and the *actual* frequency of oscillation of the presynaptic oscillator (denoted by dynamical variable *ω*\* as defined by eqns. 11 and 14). The second difference is that, in modified power coupling, the angle of the complex power coupling coefficient does not incorporate the natural frequency of the presynaptic oscillator in the denominator (conventional and modified power coupling coefficients are: 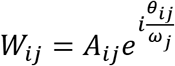 and 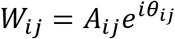 respectively). At steady state the normalized phase difference between the two oscillators 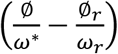 will be 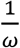 times the angle of the complex coupling coefficient (*θ_r_*) (see Appendix-1 for proof).

Under normal condition (*ε* ≠ 0) we are going to consider the special scenario where *ω*_0_ falls under the entrainment regime of the CAO oscillator. The entrainment regime of an individual oscillator, receiving only *εI*(*t*) as input, denotes the range of values of *Ω* for which the Hopf oscillator exhibits either stable fixed-point or stable spiral behavior in (*r, ψ*) space under the influence of *I*(*t*). Inside entrainment regime the actual frequency of oscillation (*ω*\*) of the oscillator is entrained to the frequency of the input signal (*ω*_0_) if the natural frequency of the Hopf oscillator (*ω*) is sufficiently close to *ω*_0_. (i.e., |*Ώ*| is sufficiently small). In fig. 3 the entrainment regime can be identified as the purple region as a function of *μ*, *β*_1_ and the intensity of the driving input (*εI*_0_). The boundary of the entrainment regime on these parameter space can also be identified as an analytical expression. From Appendix-2 we found out that the steady-state phase offset of the oscillator inside the entrainment regime is: 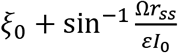. At the boundary of the entrainment regime the argument of the *arcsin* operator is 1. i.e., 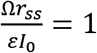, which is presented in figs. 3D, E, F as a function of one of these three parameters keeping other two fixed.

**Figure 3:**
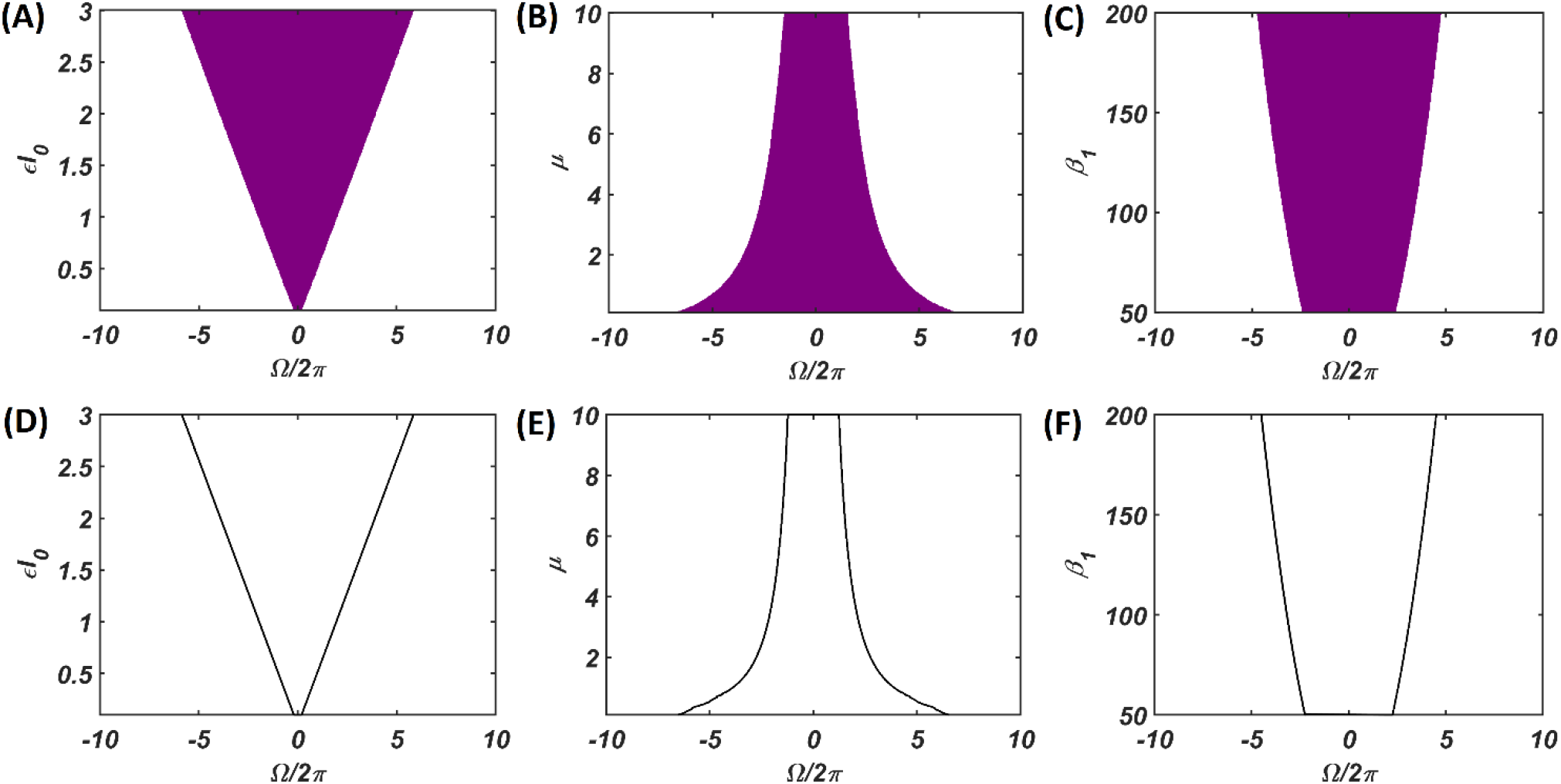
The purple region on the **(A), (B)** and **(C)** plots shows the entrainment regime of an individual Hopf oscillator and how it is dependent on its intrinsic dynamical parameters (*μ* and *β*_1_) and the intensity of the driving input (*εI*_0_). Whereas the boundaries of the entrainment regime on these parameter space plotted in **(D), (E)** and **(F)**, represented by the function 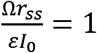. In each of these plots one of these parameters are varied while keeping others fixed at: *μ* = 1, *β*_1_ = 150, *εI*_0_ = 2.

**Figure 4:**
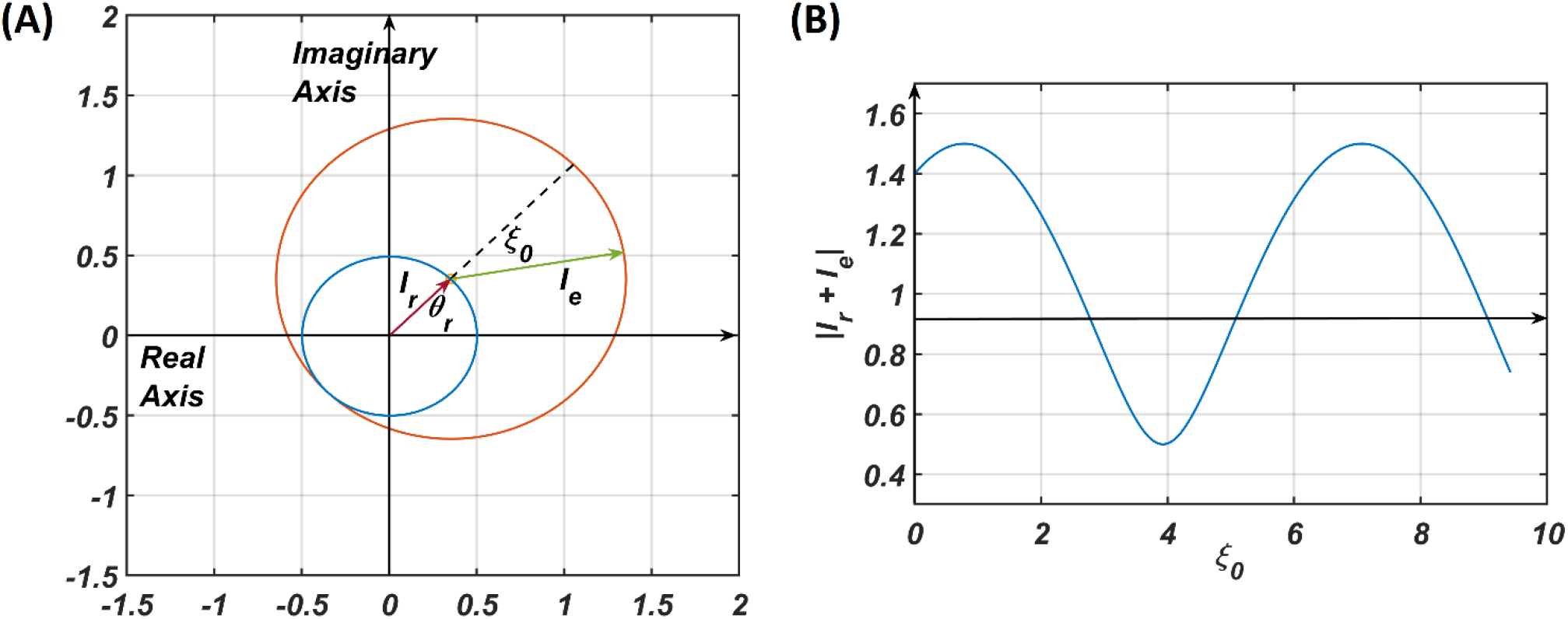
The pictorial representation of the constructive and destructive interference between the two inputs to a CAO oscillator. *I_r_* = 0.5*e^iθ_r_^*, *I_e_* = *e*^*iξ*_0_^ are the phasors of the input from the reference oscillator and the external input respectively. *I* = *I_e_* + *I_r_*. In **(B)** *θ_r_* is kept fixed at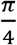 and *ξ*_0_ is varied to plot |*I*| from which it can be observed that |*I*| is maximum when *ξ*_0_ = 2*nπ* + *θ_r_* and minimum when *ξ* = (2*n* + 1)*π* + *θ_r_*.

The dependency of the width of the entrainment regime, Δ*ω* on *μ, β*_1_ and *εI*_0_ is further illustrated in fig. A2.2 in the Appendix. Although at the beginning, the CAO oscillator is perturbed by two complex sinusoidal input signals, one with frequency *ω* (from the SRO, but after the modified power-coupling step) and the other with frequency *ω*_0_ (external input *I*(*t*)), it is entrained to the external input signal because of the dominance of the perturbation by *I*(*t*) over the perturbation caused by the input from SRO since we assume that *A*_r_ < *εI*_0_. The entrainment width of the CAO oscillator will also depend on *θ_r_* – *ξ*_0_ (fig. 6). Since the SRO does not receive any inputs, it always oscillates at its natural frequency of oscillation (*ω_r_*).

Thus, under the conditions of entrainment, a given CAO oscillator gets two complex sinusoidal signals as inputs with the same frequency as *I*(*t*), with different magnitude and phase offsets, given as follows:

- 1) the 3^rd^ term in the RHS of eqn. 9 evolves to 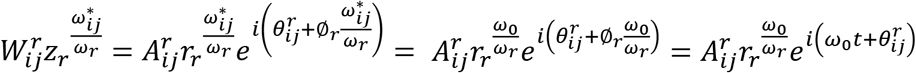 at steady state (refer to Appendix-4), denoted by *I*_r_(*t*) = *a_r_e*^*i*(*ω*_0_*t*+*ξ*_*r*_)^ and,
- 2) the 4^th^ term in RHS of eqn. 9, *εI*_0_*e*^*i*(*ω*_0_*t*+*ξ*_0_)^ denoted by *I_e_*(*t*) = *a*_0_*e*^*i*(*ω*_0_*t*+*ξ*_0_)^ (Appendix-4), where *a_r_* is dependent on the steady state magnitude of oscillation of the reference oscillator (*r_rss_*), which in turn depends on *μ_r_*, *β*_1*r*_ and *A_r_*. *ξ_r_* being *θ_r_* justifies why the actual frequency of the presynaptic oscillator is omitted from denominator of the angle of the modified power coupling coefficient. Both of these inputs either constructively or destructively interfere with each other depending on their relative phase offset (*ξ_r_* – *ξ*_0_ = *θ_r_* – *ξ*_0_). Since the magnitude of the response of the CAO oscillator is either diminished or increased, depending on the relative values of *θ_r_* and *ξ*_0_, with this arrangement the CAO oscillator will be capable of encoding phase offset of the complex sinusoidal input signal.

#### Modified Hebbian learning

A modified Hebbian learning rule is proposed for training the modified power coupling described in the previous section. The complex variable and the polar coordinate version of the modified Hebbian learning rule is described as following;

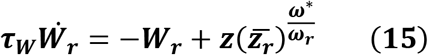

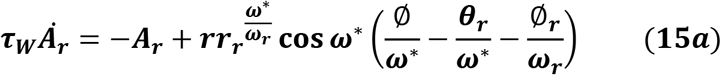

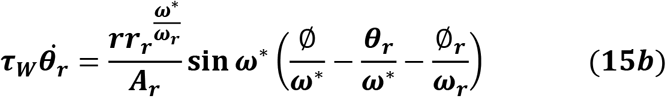

The modified Hebbian learning rule as prescribed in eqns. 15a,b can be compared to the original Hebbian learning for the power coupling coefficient proposed earlier (eqn. 15 in Biswas et al., 2021). Here 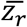 is the complex conjugate of the complex activation of the SRO. It can be observed that the modified Hebbian learning rule for the modified power coupling coefficient has a similar effect as the original Hebbian rule had in case of the previously proposed power coupling strategy.

Effectively the modified Hebbian learning rule is the same as the previous one when there is no entrainment. Without entrainment when the phase offset of the post-synaptic oscillator is driven to the phase offset of the complex sinusoidal input signal with identical frequency as the natural frequency of the CAO oscillator, *θ_r_* learns *ξ*_0_, where *ξ*_0_ is the phase offset of the input driving the post-synaptic oscillator (refer to Appendix 5). Similarly, it can be shown that *A_r_* learns 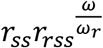 (refer to Appendix 6), where *r_ss_* and *r_rss_* are the steady state values of amplitude of oscillation of the post-synaptic and presynaptic oscillators respectively. With entrainment of the main oscillator *θ_r_* learns 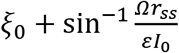, which is the phase offset of the main oscillator at steady-state after entrainment while the Hebbian dynamics is enabled.

### Training the OTSOM

In the previous section, we discussed the response properties of a single CAO oscillator to its two inputs. We now discuss the training methodology of OTSOM. The OTSOM model is trained on complex sinusoidal signals, each defined by a characteristic frequency (*ω*_0_) and phase (*ξ*_0_) (defined w.r.t. SRO). Thus, for a given sinusoidal input, we expect the OTSOM to produce a single winner, such that the row number, *i*, of the winner represents the input phase, while the column number, *j*, represents the input frequency. Therefore, the CAO oscillator at (*i,j*) gives maximum resonating response when *ω*_0_ = *ω_ij_* and 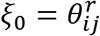 which is why we have chosen to train the *ω_ij_* and 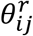 parameters of the OTSOM network. These parameters are trained in two consecutive training stages. In the 1^st^ stage, *ω_ij_*s are trained keeping 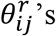 fixed and in the 2^nd^ stage, 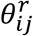s are trained by keeping the already trained *ω_ij_*s fixed.

### First Stage of training: Training Frequency

In this stage, the natural frequencies, *ω_ij_*, of the CAO oscillators are trained according to a learning rule analogous to the self-organizing map learning rule (Kohonen, 1990). Specifically, to train the frequencies of the individual oscillators, we use the adaptive frequency Hopf oscillator model proposed earlier (Biswas et al., 2021; Righetti et al., 2005). The training takes place over multiple epochs (*N_epocft,ω_*). In every epoch, *N* input signals are randomly chosen from the input set Ʊ. The input set contains complex sinusoidal signals with frequencies and phase offset sampled from a uniform probability distribution. Once an input signal with certain frequency and phase offset (*I*_0_*e*^*i*(*ω*_0*p*_*t*+*ξ*_0*p*_)^) is selected, it is presented as the external input signal (*I_p_*(*t*)) to the oscillators in the CAO. After an initial transient phase, which lasts for *T_sω_* seconds, the CAO oscillator response reaches steady state. The oscillator with the largest amplitude at the end of transient phase is denoted the ‘winner CAO oscillator’.

The dynamics of eqns. (9, 10, 11) is simulated with the special condition 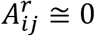 during the transient phase. Under this condition the steady state solution of CAO oscillator 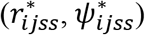 can either be a stable node, stable spiral or unstable spiral (Kim & Large, 2015). Typically, inside the entrainment regime we get stable node or stable spiral as the solution. The parameters *μ, β*_1_ and *ε* play a crucial role during this phase as they determine not only the entrainment width of the Hopf oscillator (Δ*ω*) but also the duration of the transient period (*T_sω_*). As the oscillator with the highest steady state amplitude of oscillation (max(*r_ijss_*)) is chosen to be the winner, it is evident that the winner oscillator should satisfy the condition 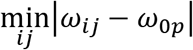. After the transient period, the frequencies *ω_ij_*s of all the CAO oscillators within the neighbourhood window, along with the winner oscillator, are trained for *T_tω_* secs according to the following dynamics (Biswas et al., 2021):

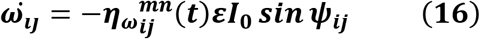

where 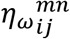 is the neighbourhood function, centered on the winner oscillator located on the *m^th^* row and column, *n^th^* defined as following:

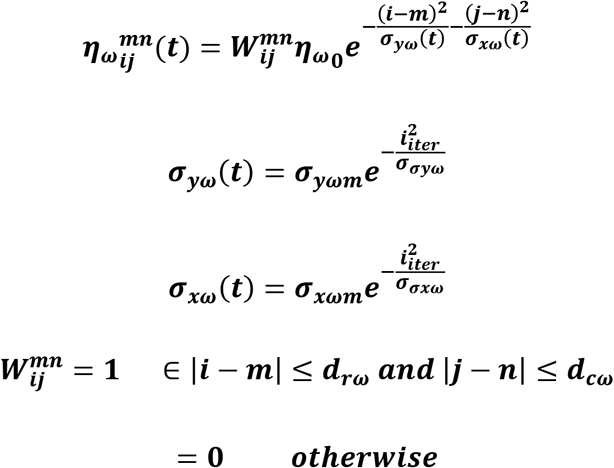

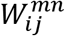 is termed as neighbourhood window. Here, *d_rω_* and *d_cω_* are the half-lengths of the neighbourhood window along rows and columns respectively. Effectively the purpose of the neighbourhood window is to constrain the learning confined to the neighbourhood of the winner oscillator. Additionally, it can be observed that the neighbourhood function, 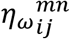, is a Gaussian centred on the winner oscillator. The standard deviations along the rows and columns (*σ_yω_* and *σ_xω_*) of 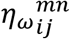 decrease with time to produce annealing effect. Thus, the network dynamics is simulated for (*T_sω_* + *T_tω_*) seconds for each presentation of a training signal, for *N*(*T_sω_* + *T_tω_*) during each training epoch, where N is the number of training signals. Therefore, the time required for the entire first phase of training is *N_epocft,ω_N*(*T_sω_* + *T_tω_*).

### Second stage training: training phase off-set

While the objective of the first stage training is to train the frequencies, *ω_ij_*s, of the CAO oscillators, the objective of the second stage training is to train the phases of the same oscillators which are determined by the angles, 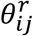, of feedforward connections from the SRO to the CAO oscillators. After the first stage training, *ω_ij_*s self-organize in an increasing order along the rows, a pattern confirmed by the simulations shown in the results section. This occurs because the CAO is a rectangular array with the number of columns much larger than the number of rows. The 2^nd^ stage of training commences with randomly initialized 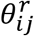 parameters and follows the algorithmic course as given in fig. 5B.

**Figure 5:**
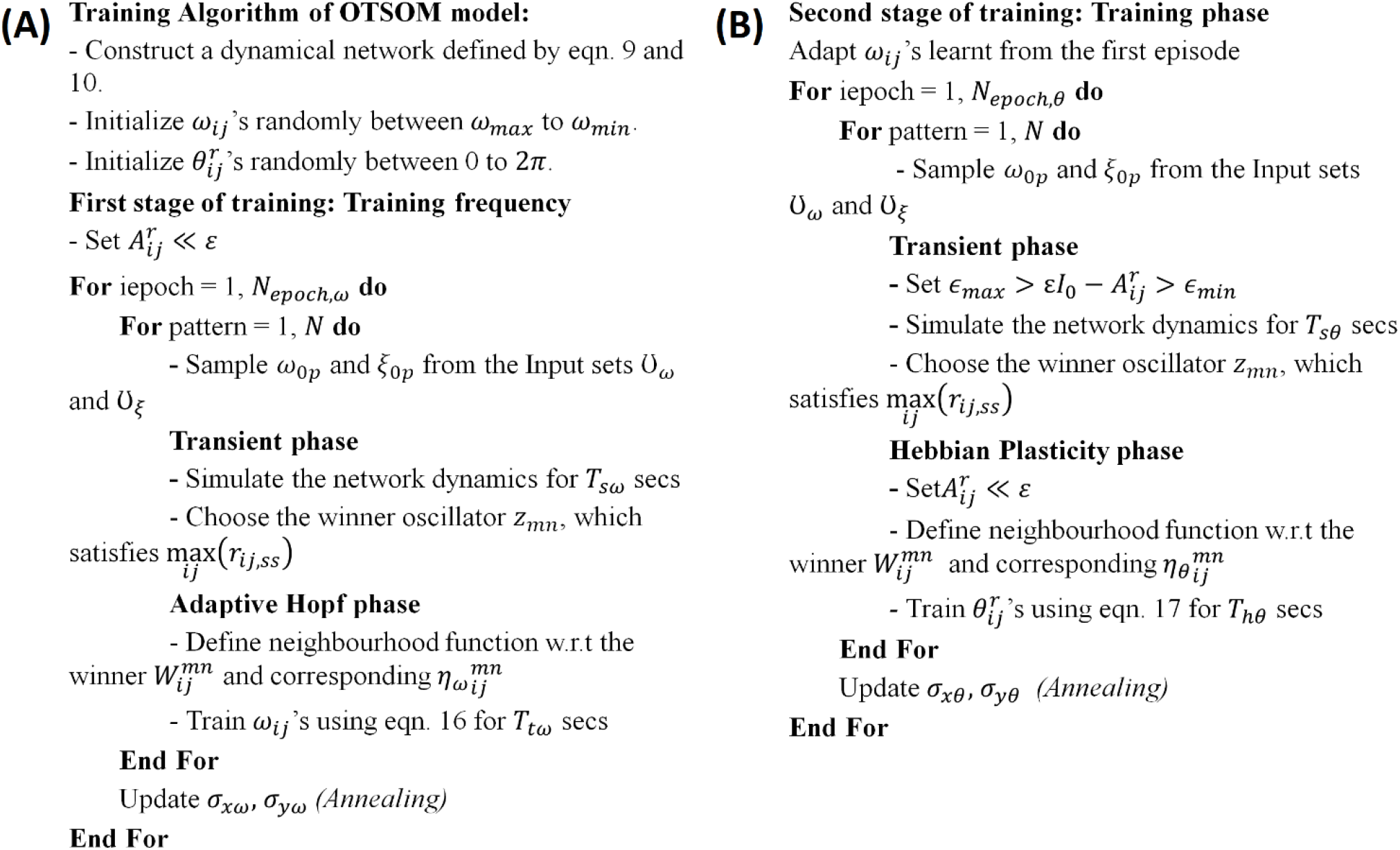
Algorithms for the first **(A)** and the second **(B)** stage of training of OTSOM model.

In the second stage training, during the transient period, the magnitudes of 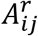 parameters are increased to the same order of magnitude as *ε*. The training takes place through multiple epochs (*N_epocft,θ_*) in which, as in the previous stage, each training pattern is a complex sinusoidal signal with specific *ω*_0*p*_ and *ξ*_0*p*_, sampled from the input set Ʊ. As in the previous training stage, the network dynamics is first allowed to reach the steady state before 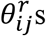 are adapted. The transient phase dynamics, given by eqns. 9, 10, 11 is simulated for *T_sθ_* secs. At the end of the transient period the winner oscillator is identified by its steady state amplitude of oscillation 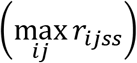. It is expected that the natural frequency (*ω_mn_*) and the angle of the power coupling coefficient 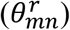 of the winner oscillator should be closest to *ω*_0*p*_ and *ξ*_0*p*_ respectively. During the subsequent *T_hθ_* period 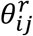 parameters of the unidirectional power coupling coefficients are trained according to the learning rule given in eqn-17. As eqn-17 is derived from the modified Hebbian learning rule as proposed in eqn-15b, this phase is termed as Hebbian learning phase.

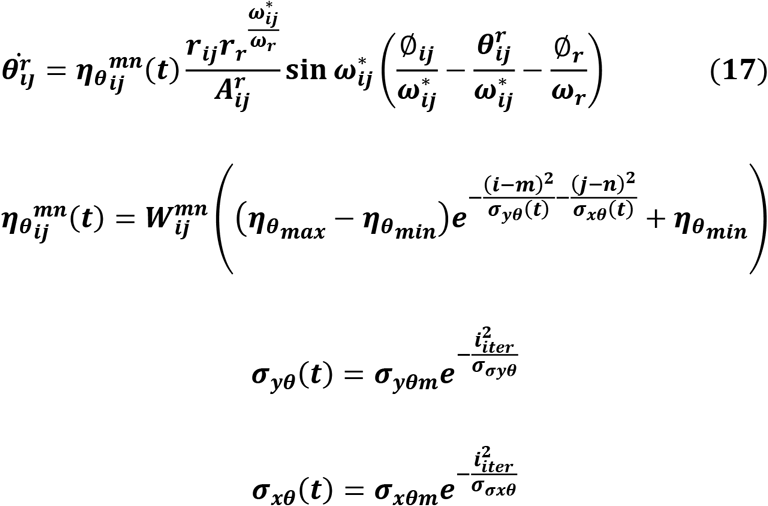

Here, *m* and *n* denote the index of the winner oscillator. The gaussian shaped neighbourhood function, 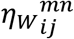, is confined between *η_Wmax_* and *η_Wmin_*. The learning neighbourhood window 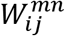 is defined w.r.t the winner oscillator similar to the first stage training. In the Hebbian learning phase 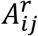 parameters are not trained 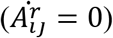, we have kept 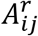 values close to zero or 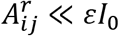. In which case, 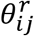 learns 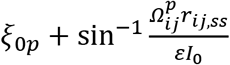 at steady-state when the CAO oscillator receives the input *I*_0_*e*^*i*(*ω*_0*p*_*t*+*ξ*_0*p*_)^ (refer to Appendix-5) (where 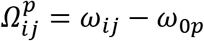). It has also been shown that even if 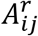 is not negligible (i.e., 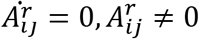) 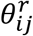 learns 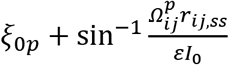 at steady-state (refer to Appendix-6). This happens because the phase offset of the CAO oscillator (*δ_ij_*) attains the value 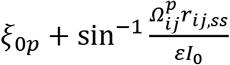 at steady state inside the entrainment regime. Each training epoch in the 2^nd^ stage training takes *N*(*T_sθ_* + *T_hθ_*) seconds, and the time required for the entire 2^nd^ stage of training is *N_epocft,θ_ N*(*T_sθ_* + *T_hθ_*).

## 3 Results

### Single oscillator results

We now numerically analyse the response of a single CAO oscillator to the simultaneously received inputs from SRO (*I_r_*(*t*)) and the external input (*I_e_*(*t*)) for the following conditions:

a. *A_r_* being sufficiently smaller but not negligible w.r.t *εI*_0_ (i.e., the strength of the *I_e_*(*t*) combining its own magnitude of oscillation and the afferent weight) which enables the CAO oscillator to get entrained to *I*(*t*), and ensures significant interference between the two inputs. The condition can be mathematically rephrased as:

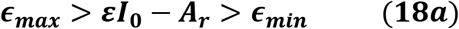

where *ϵ_min_* and *ϵ_max_* are positive numbers.
b. *A_r_* is negligible w.r.t *εI*_0_, which is similar to the scenario where CAO oscillator is only perturbed by *I*(*t*).

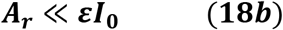

We have numerically analyzed the response of the CAO oscillator under the 1^st^ condition (eqn. 18a) as described by the eqns.- 9,10,11. The CAO oscillator exhibits stable entrainment depending on both the relative frequency (*Ω*) and the relative phase (*θ_r_* – *ξ*_0_) w.r.t the external input signal. When it shows stable entrainment, the magnitude of oscillation reaches a fixed value at steady state. There is a distinguishable boundary on the *Ω* vs *θ_r_* – *ξ*_0_ plane in which stable entrainment is observed. Outside this boundary the CAO either exhibits intermittent entrainment or does not get entrained at all. In the following figure (fig. 6) the region in the *Ω* vs *θ_r_* – *ξ*_0_ plane where stable entrainment is observed is portrayed as purple region. It can be observed that there is a symmetry in the purple region w.r.t the *Ω* = 0 and *θ_r_* – *ξ*_0_ = 0 axis (*θ_r_* = *π*). A pair of CAO oscillator and a SRO is simulated until the steady state is achieved to find out the mean, maximum and the minimum value of the steady state magnitude of oscillation of the CAO oscillator. The following parameters are used: *μ* = 1, *β*_1_ = –100, *ω* = 2*π*60, *ω*_0_ = 2*π*50 to 2*π*70, *ξ*_0_ = 0 to 2*π, ε* = 2, *F* = 1, *μ_r_* = 1, *β*_1*r*_ = –10, *ω_r_* = 2*π* × 60, *τ_ω_* = 0.1, *A_r_* = 0.5, *θ_r_* = *π* for the simulation results provided in the figures (figs. 7,8,9). The steady state is attained typically before 3 secs. The steady state behaviour of the magnitude of oscillation of the CAO oscillator for eight different symmetrical cross sections parallel to *Ω* = 0 axis and for another eight different symmetrical cross sections parallel to *θ_r_* – *ξ*_0_ = 0 axis are plotted in the fig. 7 and fig. 8.

**Figure 6:**
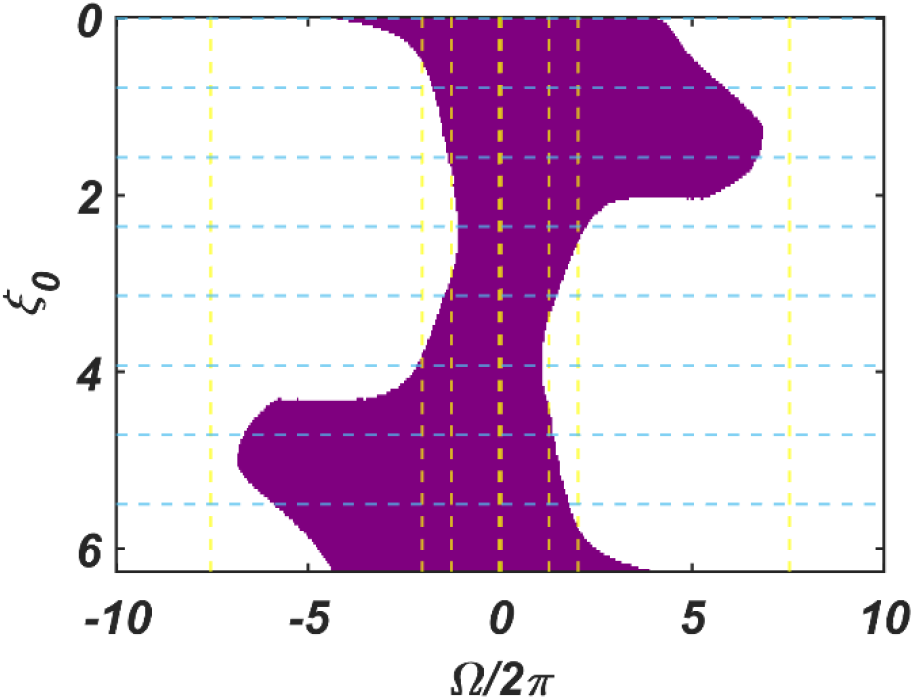
The figure principally depicts the entrainment regime (purple region) of the CAO oscillator on the *Ω* vs *θ_r_* – *ξ*_0_ plane, keeping the other factors such as *εI*_0_, *θ_r_, A_r_* and the intrinsic parameters of the CAO oscillator and the SRO fixed.

**Figure 7:**
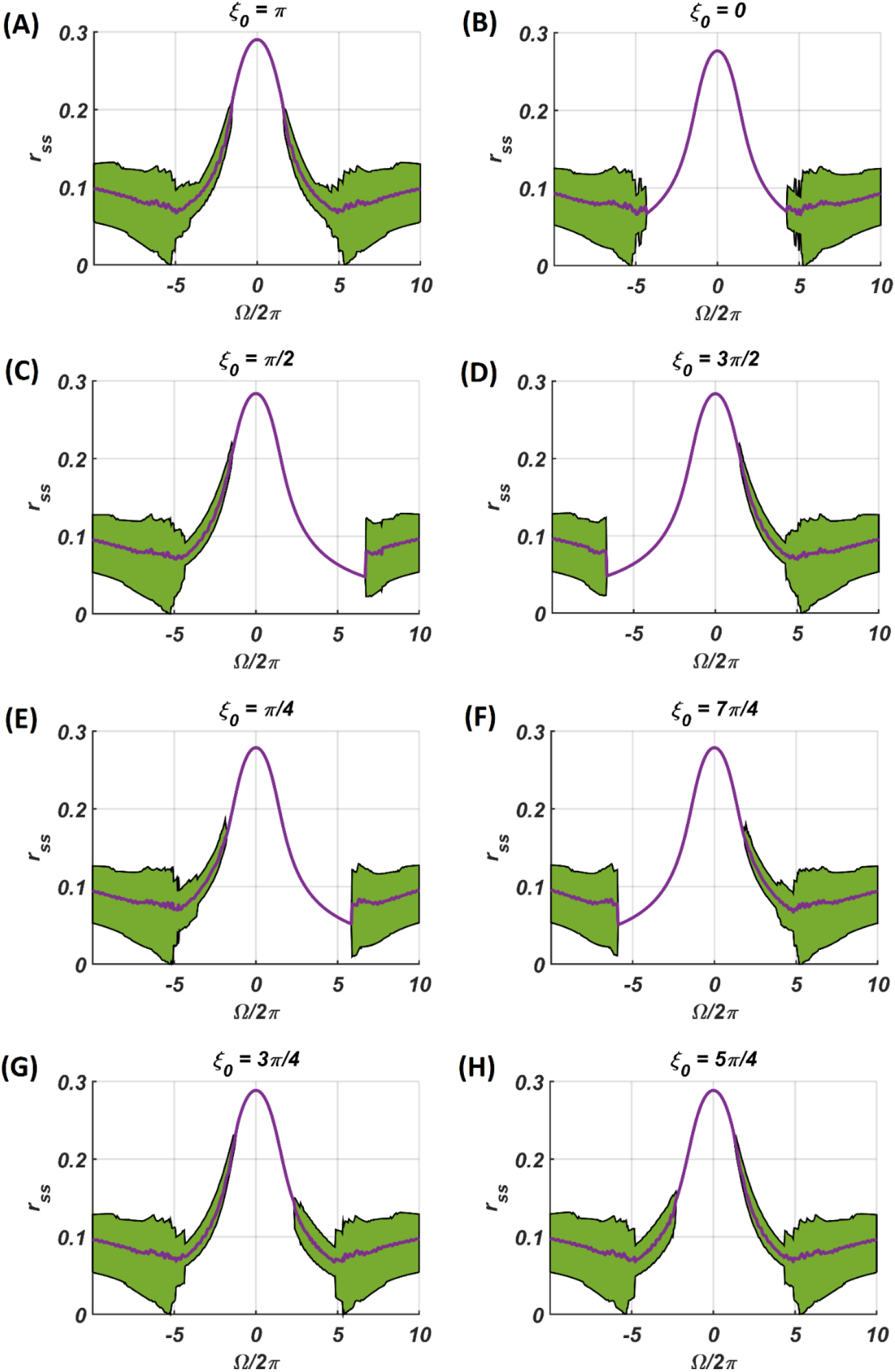
The steady-state magnitude of oscillation of the CAO oscillator with the identical pair of coupled CAO oscillator and SRO as provided in the fig. 6 but for a symmetrical distinct value of the phase offset of the external input signal w.r.t the angle of the power coupling connection *θ_r_* = *π*, cross section of which is drawn in blue lines in fig. 6. The green region depicts the variance of steady state magnitude of oscillation w.r.t the mean value, plotted in purple.

**Figure 8:**
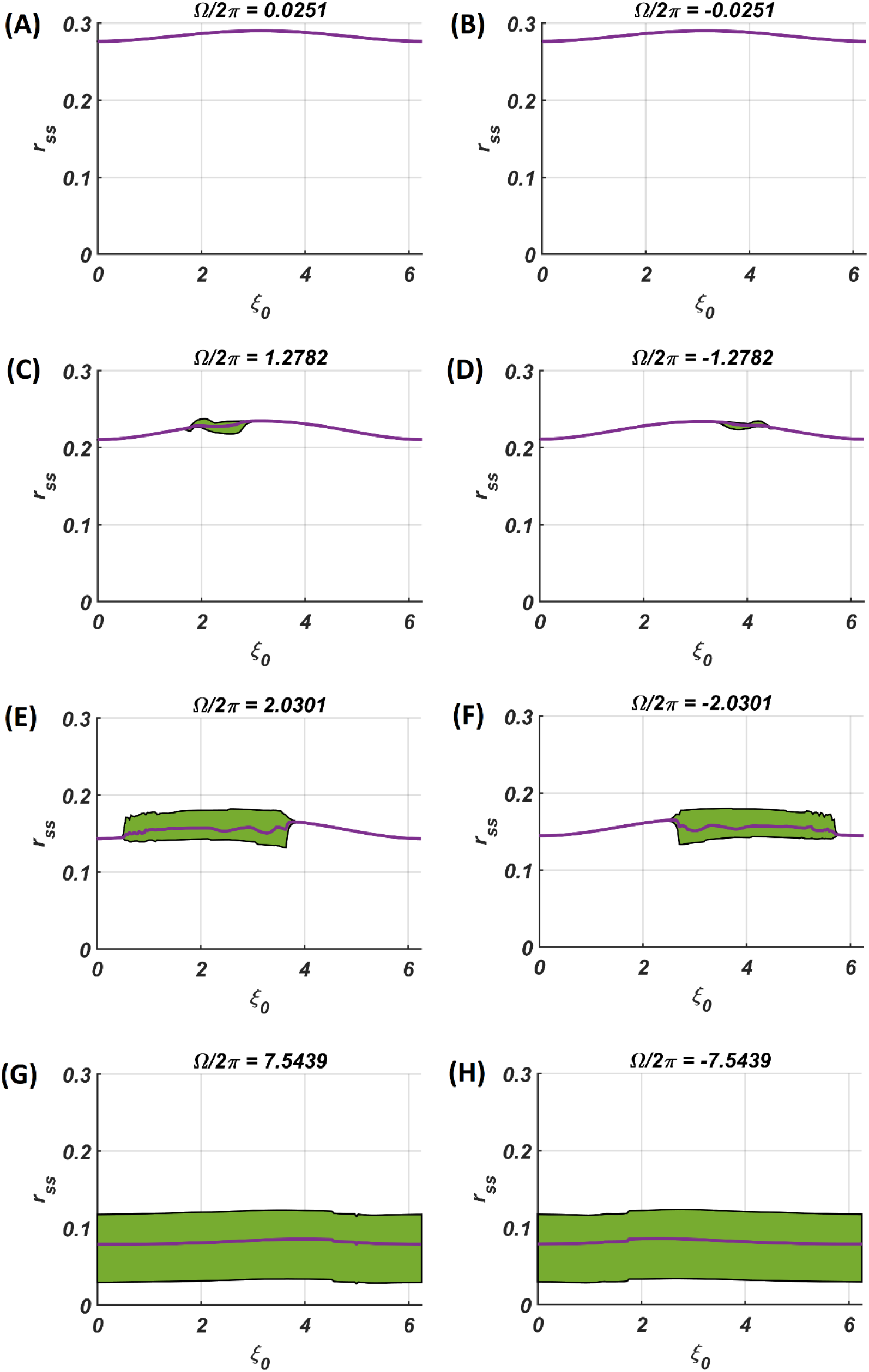
The steady-state magnitude of oscillation of the CAO oscillator with the identical pair of coupled CAO oscillator and SRO as provided in the fig. 6 but for a symmetrical distinct value of the relative frequencies of the external input signal w.r.t the natural frequency of the CAO oscillator *ω* = 2*π* × 60, cross section of which is drawn in yellow lines in fig. 6. The green region depicts the variance of steady state magnitude of oscillation w.r.t the mean value, plotted in purple.

**Figure 9:**
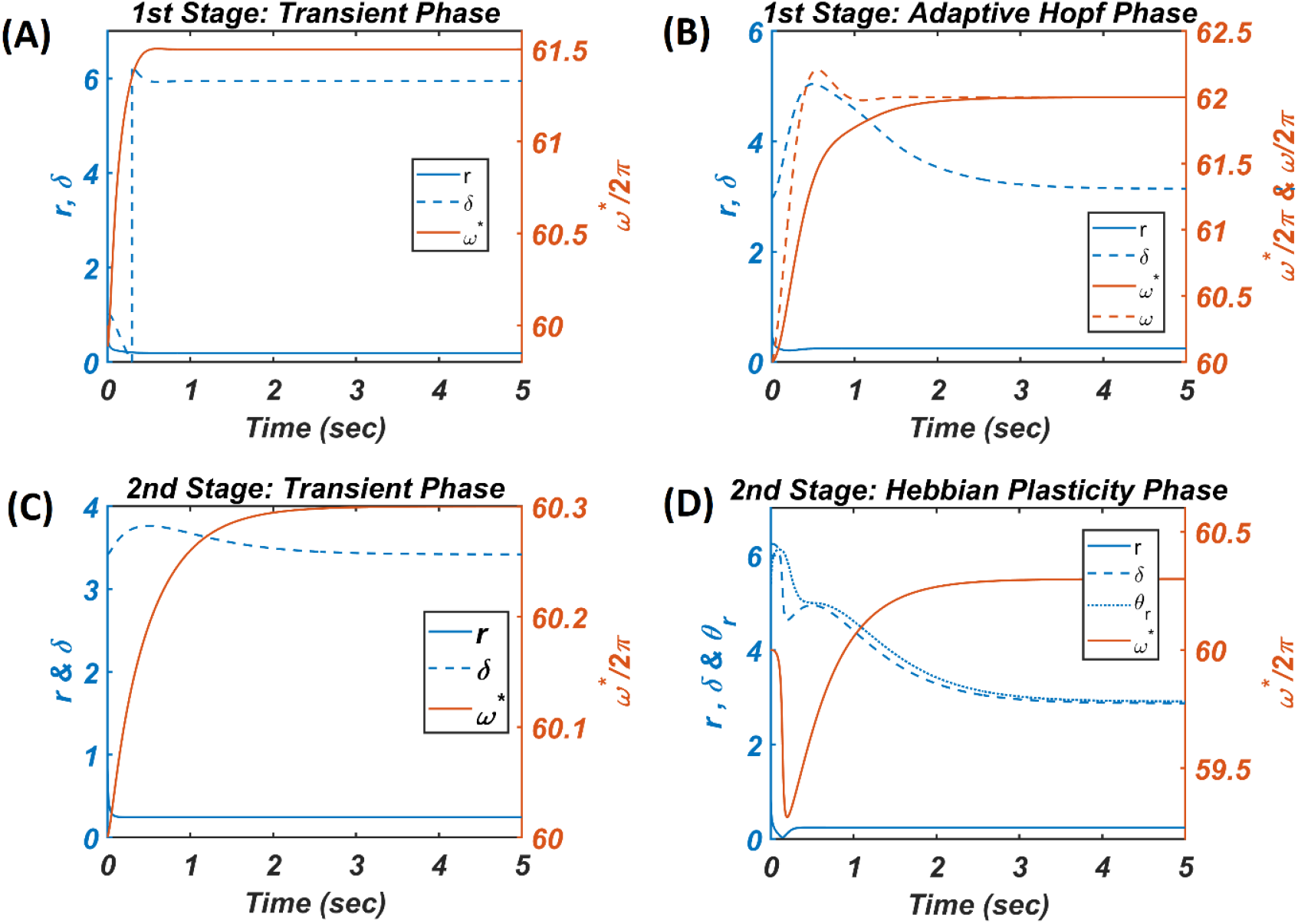
The figure depicts the response of the CAO oscillator in terms of its magnitude of oscillation (*r*), phase offset (*δ*), entrainment (*ω*\*) during all the four phases of the 1^st^ and 2^nd^ stages of training along with the parameters to be trained (*ω* and *θ_r_*) and the other fixed parameters *μ* = 1, *β*_1_ = 150, *ε* = 2, *I*_0_ = 1, *μ_r_* = 1, *β*_1*r*_ = 10, *ω_r_* = 2*π* × 60.5. **(A)** During the transient phase of the 1^st^ stage of training the single unit is simulated under the entrainment regime of the CAO oscillator with the additional parameters: *ω*_0_ = 2*π* × 61.5, 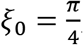, *τ_ω_* = 8 × 10^-4^, *A_r_* = 10^-5^. It can be verified that at steady-state *r* and *δ* attain the solution provided by eqn. A1.8 and the value 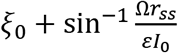 respectively. **(B)** For the adaptive Hopf phase of the 1^st^ stage of training the single unit is simulated with the additional parameters: *ω*(0) = 2*π* × 60, *ω*_0_ = 2*π* × 62, *ξ*_0_ = *π*, *θ_r_* = *π, τ_ω_* = 0.5, = 10^-5^, *μ_ω_* = 50. It can be verified that at steady state *η* attains *ξ*_0_ and *ω* learns *ω*_0_. **(C)** The single unit is simulated with the following set of additional parameters during the transient phase of the 2^nd^ stage of training: *ω*_0_ = 2*π* × 60.3, *ξ*_0_ = 3.6773, *τ_ω_* = 0.5, = 0.1. The steady-state values of *r* depends on 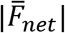 whereas the steady-state value of *δ* is same as 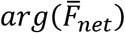. **(D)** For the Hebbian plasticity phase of 2^nd^ stage of training the single unit is simulated with the additional parameters: *ω*_0_ = 2*π* × 60.3, *ξ*_0_ = *π*, *τ_ω_* = 0.5, *A_r_* = 10^-5^, *μ_θ_* = 10^-5^. It can be verified that *θ_r_* learns 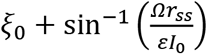 which is same as the steady-state value of *δ*.

In the figs. 8 and 9 the *r_ss_* is plotted on a same scale of magnitude so that the resonance exhibited by the CAO oscillator w.r.t the frequency and the phase offset of the external input can be compared. It is apparent that *ω* has a greater effect on CAO oscillator in terms of its steady-state magnitude of oscillation than *ξ*_0_. The reason behind introducing the lower bound on *εI*_0_ – *A_r_*(*ϵ_min_*) is justified here. In other words, the resonance exhibited by the CAO oscillator w.r.t the *ξ*_0_ at a smaller scale is ensured by the condition *εI*_0_ – *A_r_* > *ϵ_min_*. The combined results of figs. 7, 8, 9 reveals that the CAO oscillator will oscillate with maximum magnitude at steady-state when |*Ω*| × |*θ_r_* – *ξ*_0_| is minimum.

Considering that the CAO oscillator is operating in the entrainment regime under the 1^st^ condition (eqn. 18a) and gets entrained to *I*(*t*), the magnitude and the phase offset of oscillation at steady state is analytically derived in Appendix-4. As the Hopf oscillator always maintains the same phase offset as the phase offset of the complex sinusoidal input signal, the phase offset of the CAO oscillator becomes the phase offset of the resultant input 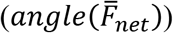, obtained by combining the external input, and the input from the SRO through the complex power coupling connection (elaborated in Appendix-4).

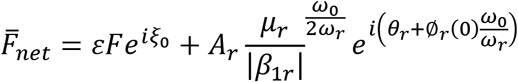

The first term on the right-hand side is the phasor of the external input (*I*(*t*)) through afferent weight, and the second term is the phasor of the input from the SRO through modified power coupling. Note that the magnitude of the resultant input after constructive/destructive interference, 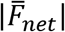, determines the magnitude of oscillation of the CAO oscillator at steady state. Fig-10C numerically verifies that the CAO oscillator attains the analytically derived solutions in Appendix-4.

**Figure 10:**
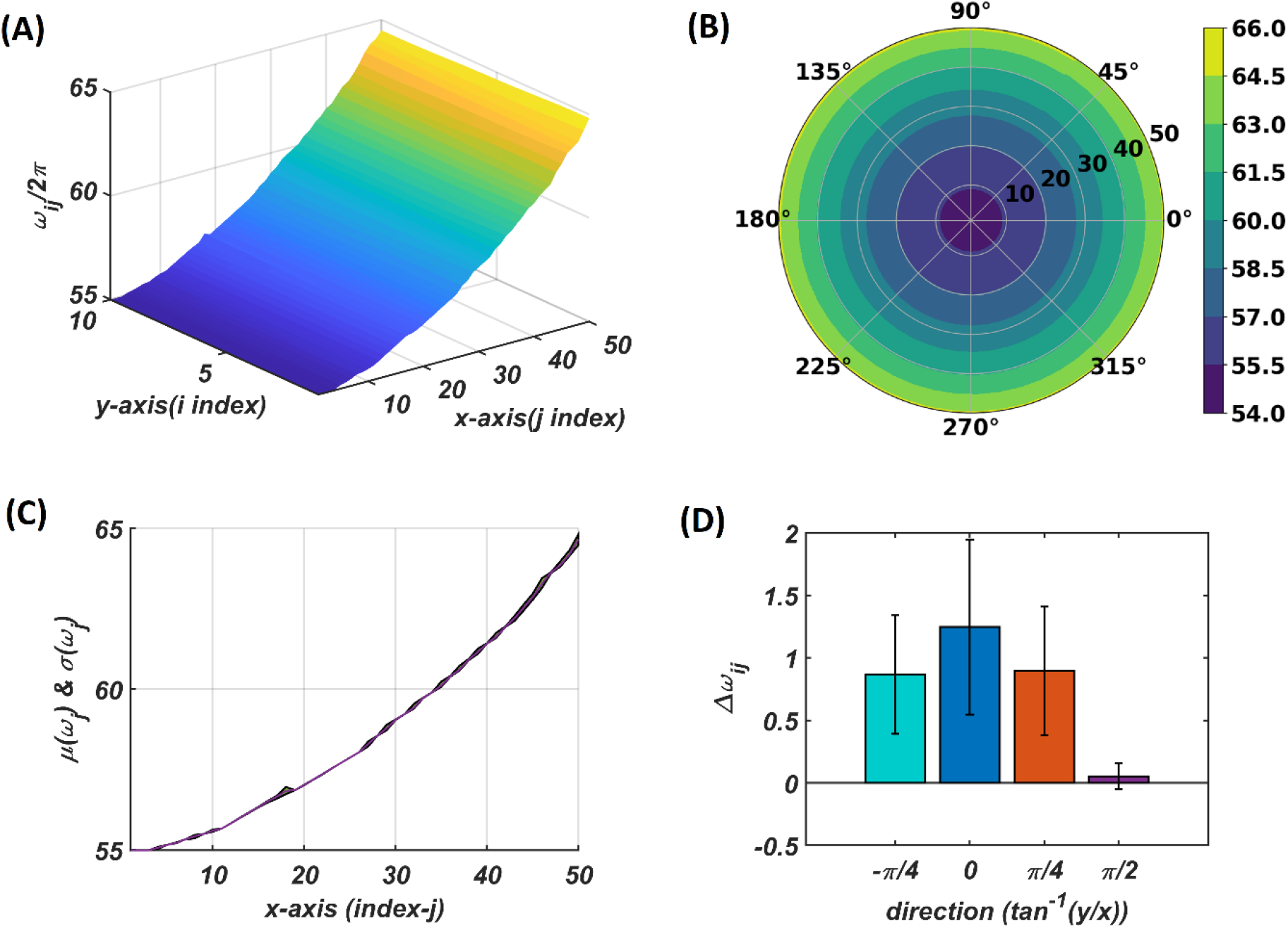
The natural frequency of oscillation, *ω_ij_*, of the oscillators in the CAO after self-organization represented in Cartesian **(A)** and polar **(B)** organization. **(C)** shows mean and the variance of the natural frequencies along x-axis after training. **(D)** provides a quantitative understanding of the slope of self-organized *ω_ij_*s along the x-axis, y-axis, and the axes 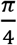 and 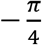 inclined w.r.t the x-axis.

The effect of *μ, β*_1_ and *εI*_0_ on the entrainment width (Δ*ω*) and the typical transient time (*T_t_*) of the CAO oscillator for the 2^nd^ condition (eqn.18b), is further verified numerically. From fig. A2.2 in Appendix-2 it can be verified that *μ* has to be small for wider entrainment regime, i.e., higher values of *μ* causes shrinking of the entrainment regime. On the contrary *ε* and *β*_1_ have nearly the same effect on the width of the entrainment regime: entrainment regime broadens as *ε* and *β*_1_ values are increased. Since the transient dynamics determines the amount of time the network takes to settle down so that the oscillator with the highest resonant response can be chosen, it is necessary to understand the effect of these parameters on the transient dynamics. It is quite intuitive that *μ* and *β*_1_ have a shrinking effect on the transient period as it causes a much steeper basin of attraction around the steady state solution. Figs. A2.2D and A2.2F show that increasing the magnitude of the scalar afferent weight, *ε*, shortens the transient period; this is natural because a stronger input pushes the oscillator output to settle down faster. Ideally, the model requires a broad entrainment regime with a short transient period.

### Network level results

#### First stage of training: training frequency

We may recall from the previous section that the objective of the 1^st^ stage of training is to train the frequencies of the CAO oscillators. To this end, only the external input signals are considered, and the inputs from SRO are ignored (i.e., *A_r_* ≪ *ε* in eqns. 9, 10, 11). Initially the input set Ʊ is constructed by sampling *N_f_* number of intrinsic frequencies 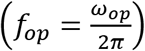 from a uniform distribution over the range of 55 to 65 Hz, and sampling *N_ξ_* number of phase offset angles (*ξ*_0*p*_) from a uniform distribution over [0, 2*π*). Combining these sampled intrinsic frequencies and phase offsets, *N* = × *N_f_* × *N_ξ_* number of complex sinusoidal signal patterns (*I_p_*(*t*) = *I*_0_*e*^*i*(*ω*_0*p*_t+*ξ*_0*p*_)^) are generated. After *I_p_*(*t*) chosen randomly from Ʊ, the input patterns are presented one at a time to all the oscillators in CAO. Note that, in this case, the winning CAO oscillator depends only on the condition 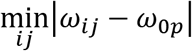, and not on the phase offset, *ξ*_0*p*_, since *A_r_* ≪ *εI*_0_.

For each presentation of the input, the winner oscillator and the oscillators in its neighbourhood, adjust their ‘preferred frequency’ closer to the input frequency, *ω*_0*p*_, following the adaptive Hopf learning rule of eqn. 16. The preferred frequency of the oscillators in the neighbourhood of the winner oscillator gets closer to *ω*_0*p*_ compared to the oscillators near the periphery of the adaptive neighbourhood because of the Gaussian neighbourhood function, 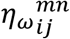. The transient response of the CAO oscillator of a single unit in terms of its magnitude of oscillation (*r*), phase offset (*δ*), entrainment (*ω*\*) during the transient phase is plotted in fig. 9A, where the analytically derived solutions of *r* and *δ* (refer to Appendix-2) is verified numerically. Similarly, the dynamic response of the same CAO oscillator in the given single unit in terms of *r, δ, ω*\* and *ω* during the adaptive Hopf phase is presented in fig. 9B, where the analytically derived solutions of *r, δ* and *ω* (refer to Appendix-3) is verified numerically.

The natural frequencies of the CAO oscillators are initialized from a uniform random distribution confined to the interval 2*π*[55,65]. The natural frequency of the reference oscillator is set to the central frequency (2*π*60) of the given frequency band so that there is a symmetrical influence by the reference oscillator on CAO oscillators. The size of the CAO (*N_r_,N_c_*) = (10, 50), where *N_r_* is the number of rows and *N_c_* is the number of columns. The adaptive neighbourhood is defined by immediate proximity rather than the physical proximity, i.e., a neighbourhood size of *d_W_* = 2 means the immediate 2 oscillators on every side of the winner oscillator including the oscillators situated diagonally.

The tonotopic arrangement emerges at the end of the 1^st^ stage of training. It can be observed from fig. 10 that *ω_ij_*s organize themselves in an increasing order along *x*-direction or increasing column index but remains almost invariant along the orthogonal (row) direction. This emergent tonotopic organization is the key feature of a self-organizing map. The parameter values defining the network architecture, the data set and the training are given in Table-1 and Table-2.

**Table-1:** The table summarizes the list of parameters defining the overall network architecture and the intrinsic parameters as well as the defining parameters of the training data set.

| NETWORK PARAMETERS | VALUES | TRAINING SET | VALUES |
| --- | --- | --- | --- |
| $N_x$ | 50 | $U_\omega$ | $\omega_{op} \sim U(55 \times 2\pi, 65 \times 2\pi)$ |
| $N_y$ | 10 | $U_\xi$ | $\xi_{op} \sim U(0, 2\pi)$ |
| $\varepsilon$ | 2 | $N_\omega$ | 60 |
| dt | 0.1 ms | $N_\xi$ | 10 |
| $A_{ij}^r$ | $10^{-5}$ | $N = N_\omega N_\xi$ | 600 |
| $\omega_r$ | $60 \times 2\pi$ | | |
| $\mu$ | 1 | | |
| $\mu_r$ | 1 | | |
| $\beta_1$ | 150 | | |
| $\beta_{1r}$ | 10 | | |

**Table-2:** The table summarizes the list of parameters essential for the 1^st^ and the 2^nd^ stage of training.

**Table-1:** The table summarizes the list of parameters defining the overall network architecture and the intrinsic parameters as well as the defining parameters of the training data set.
| 1<sup>st</sup> STAGE PARAMETERS | VALUES | 2<sup>nd</sup> STAGE PARAMETERS | VALUES |
| --- | --- | --- | --- |
| $n_{iter,\omega}$ | 500 | $n_{iter,\xi}$ | 300 |
| $T_{s\omega}$ | 3 sec | $T_{s\theta}$ | 3 |
| $T_{t\omega}$ | 4 sec | $T_{t\theta}$ | 4 |
| $\eta_{\omega_0}$ | 0.2 | $\eta_{\theta_{max}}, \eta_{\theta_{min}}$ | $2 \times 10^{-6}, 10^{-7}$ |
| $\sigma_{y\omega m}$ | 100 | $\sigma_{y\theta m}$ | 2 |
| $\sigma_{x\omega m}$ | 4 | $\sigma_{x\theta m}$ | 2 |
| $\sigma_{\sigma x \omega}$ | $500n_{eph, \omega}$ | $\sigma_{\sigma x \theta}$ | ∞ |
| $\sigma_{\sigma y \omega}$ | $500n_{eph, \omega}$ | $\sigma_{\sigma y \theta}$ | ∞ |
| $d_r$ | 3 | $d_r$ | 3 |
| $d_c$ | 3 | $d_c$ | 3 |
| $A_{ij}^r$ | $10^{-5}$ | $A_{ij}^r$ | 0.1 in transient phase and $10^{-5}$ in Hebbian plasticity phase |

#### Second stage of training: training phase offset

As described in fig. 5B the 2^nd^ stage of training commences with the tonotopically organized *ω_ij_*’s and seeks to train 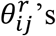. The frequency and the phase offset of the external input are sampled from the same set Ʊ. During the transient phase 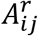 is typically set to about a tenth of *ε* so that the entrainment to *I_e_*(*t*) can occur. From fig. 9C it can be observed that entrainment is possible even if *A_r_* is comparable with *ε*. It can be assumed that the winner oscillator will be the one which is going to satisfy the condition 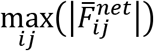, where 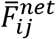 is the resultant input to the oscillator at (*i,j*) in CAO. 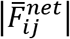 is a result of constructive or destructive interference, between the external input and the input from the SRO, depending on the relative values of *ξ*_0_ and *θ_r_*, a qualitative explanation of which is discussed in methods section with fig. 4 and the analytical expression of which is presented by eqns. A4.3 and A4.4 in Appendix-4.

The duration of the transient phase (*T_sθ_*) in the 2^nd^ stage, is typically the same as the duration of the transient phase (*T_sω_*) of the 1^st^ stage of training. With the proper choice of the network parameters, principally *ε*, 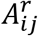, *μ, μ_r_, β*_1_, *β*_1*r*_, as given in Table-1 and 2, the duration of the transient period (*T_sθ_*) turned out to be approximately 3 secs. The CAO oscillator whose parameters 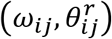 are closest to the input signal parameters (*ω*_0*p*_, *θ*_0*p*_) will be the winner.

In the following Hebbian learning phase the neighbourhood size (*d_r_* × *d_c_*) is dependent on the size of the entrainment window as the Hopf oscillator can only have a stable phase offset when it operates inside its entrainment regime. From fig. 9D it can be observed that when the angle of the complex power coupling coefficient 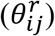 is trained according to the complex Hebbian learning rule (eqn. 17) under the 2^nd^ condition (eqn. 18b) and 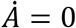, it learns the phase offset of the CAO oscillator. The transient response of the CAO oscillator of a single unit in terms of its magnitude of oscillation (*r*), phase offset (*δ*), entrainment (*ω*\*) during the transient phase is plotted in fig. 9C, where the analytically derived solutions of *r* and *δ* (refer to Appendix-4) is verified numerically. Similarly, the dynamic response of the same CAO oscillator along with the angle of the complex modified power coupling coefficient in the given single unit in terms of *r, δ, ω*\* and *θ_r_* during the Hebbian plasticity phase is presented in fig. 9D, where the analytically derived solutions of *r, δ* and *ω* (refer to Appendix-3) is verified numerically. As *ω_ij_* parameters are self-organised in a monotonically increasing fashion along the x-axis and remains almost invariant along the other orthogonal dimension, the span of the trainable neighbourhood window along the x-axis has to be chosen dependent on the entrainment width of the cortical oscillators.

With the parameters in Table 2, from fig. 11 it can be observed that the 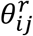 parameters have self-organized with a linear gradient along the y-axis but almost invariant along the x-axis. Fig. 11 represents four different instances of 2^nd^ stage of training with the pre-trained *ω_ij_* parameters after the 1^st^ stage of training as presented in fig. 10. The common features about all these four instances are: 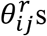 have either self-organised themselves in a linearly increasing or a decreasing fashion along the y-axis or the azimuth direction in the polar coordinate system representation, it is unpredictable where exactly along the x-axis the self-organized 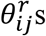 switch from an increasing to decreasing fashion and vice-versa, the overall gradients of the self-organized 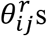 along the 4 axes (x, y and the axes inclined at an angle 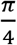 and 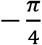 w.r.t the x-axis) for all these four instances are statistically similar. If the Gaussian shape of the neighbourhood function of 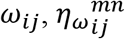, defined by its standard deviations along x- and y-axis (*σ_xω_, σ_yω_*) is compared with the Gaussian shape of the neighbourhood function of 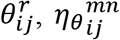, (*σ_xθ_, σ_yθ_*), the variances of 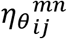 is much smaller than the variances of 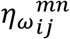, particularly along the y-axis. Also, the distribution of 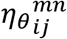has a similar spread along both of the orthogonal axes. Distribution of *ω_ij_s* from fig. 10D compared to the distributions of 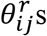 presented in fig. 11B, D, E and F confirms that they self-organize along orthogonal directions.

**Figure 11:**
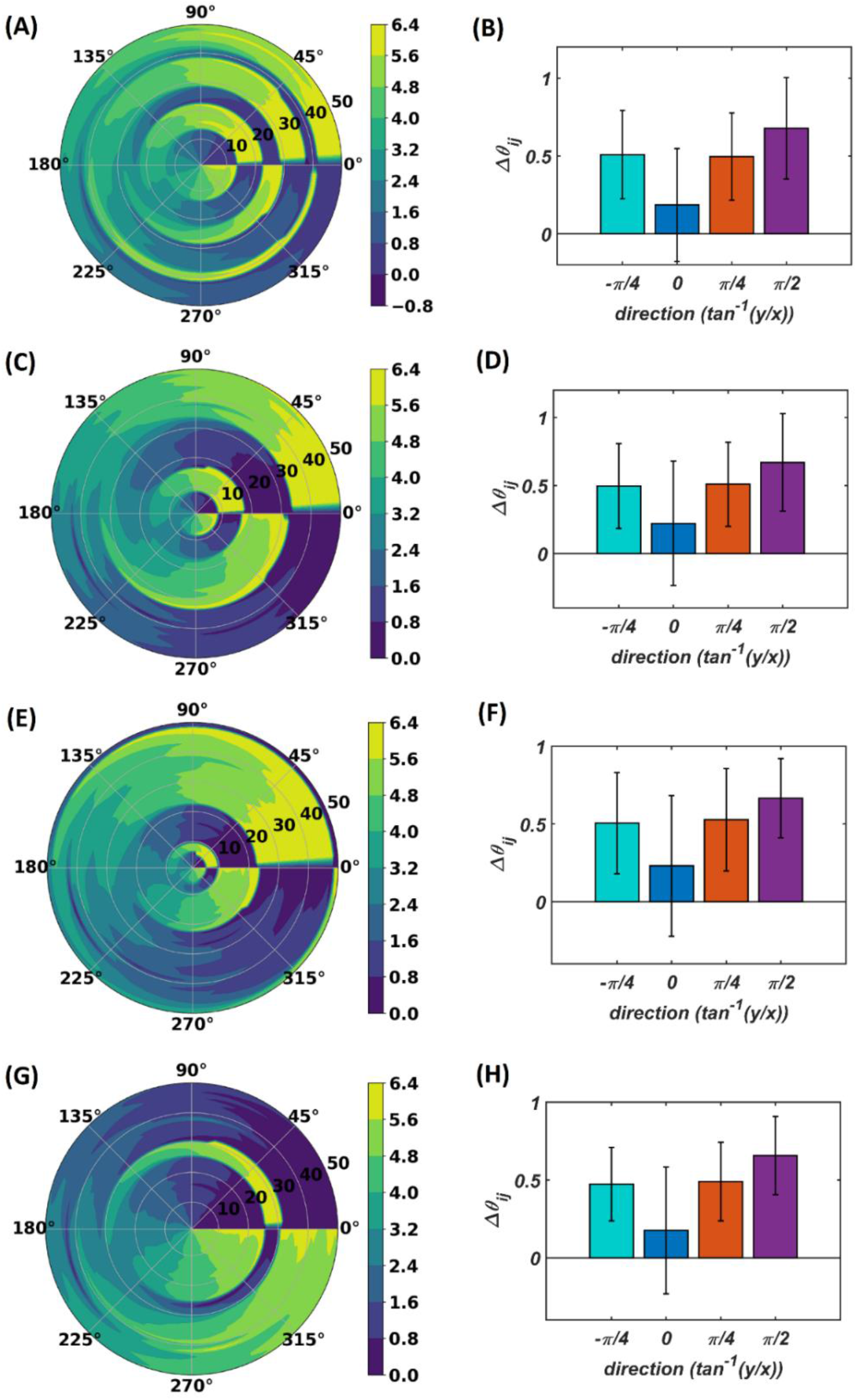
The self-organized 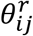 parameters after the 2^nd^ stage of training on four separate instances starting with different randomly initialized 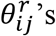. The subplots **(B)**, **(D)**, **(F)** and **(H)** represent the mean and the standard deviation of the gradients along the radial (0), azimuth 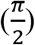, spirally diverging 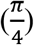 and converging 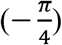 direction on a polar coordinate representation of the self-organized 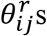 presented in the subplots **(A)**, **(C)**, **(E)** and **(F)** respectively after the 2^nd^ stage of training.

### Testing

During the testing phase of the OTSOM model, the conventional power coupling is used from the SRO to the CAO oscillator. The dynamics of the model during the testing phase is described as below:

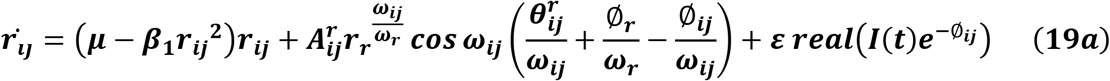

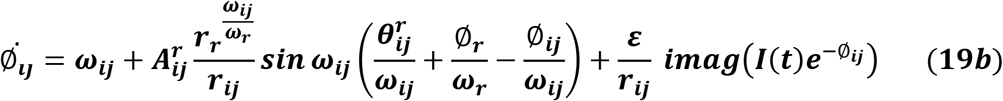

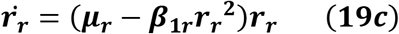

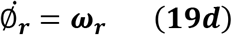

It can be observed that the ratio of the natural frequencies of the CAO oscillators w.r.t the reference oscillator is raised to power of the complex activation of the reference oscillator instead of the actual frequencies of the CAO oscillator w.r.t the reference oscillator. For the moment, real sinusoidal signals or linear combination of multiple real sinusoidal signals are used as external input (*I*(*t*)). At first, we are going to analyse the steady state response of a single CAO oscillator by considering the DC component of the magnitude of oscillation of the CAO oscillator numerically. The DC component of the steady state magnitude of oscillation is extracted using a third order low pass filter with a cut off frequency of 0.01 Hz. From fig. 12 it can be observed that the DC component of the steady state magnitude of oscillation preserves the encoding ability of all the characteristic components of the sinusoidal input signal. The resonance exhibited w.r.t the frequency of real sinusoidal external input is similar to the case of complex sinusoidal external input as observed in figs. 13A and D. Whereas the resonance exhibited w.r.t the phase offset of real sinusoidal external input is confined to a very narrow band-width of the frequency of real sinusoidal external input w.r.t the natural frequency of the CAO oscillator (figs. 13B, 13E).

**Figure 12:**
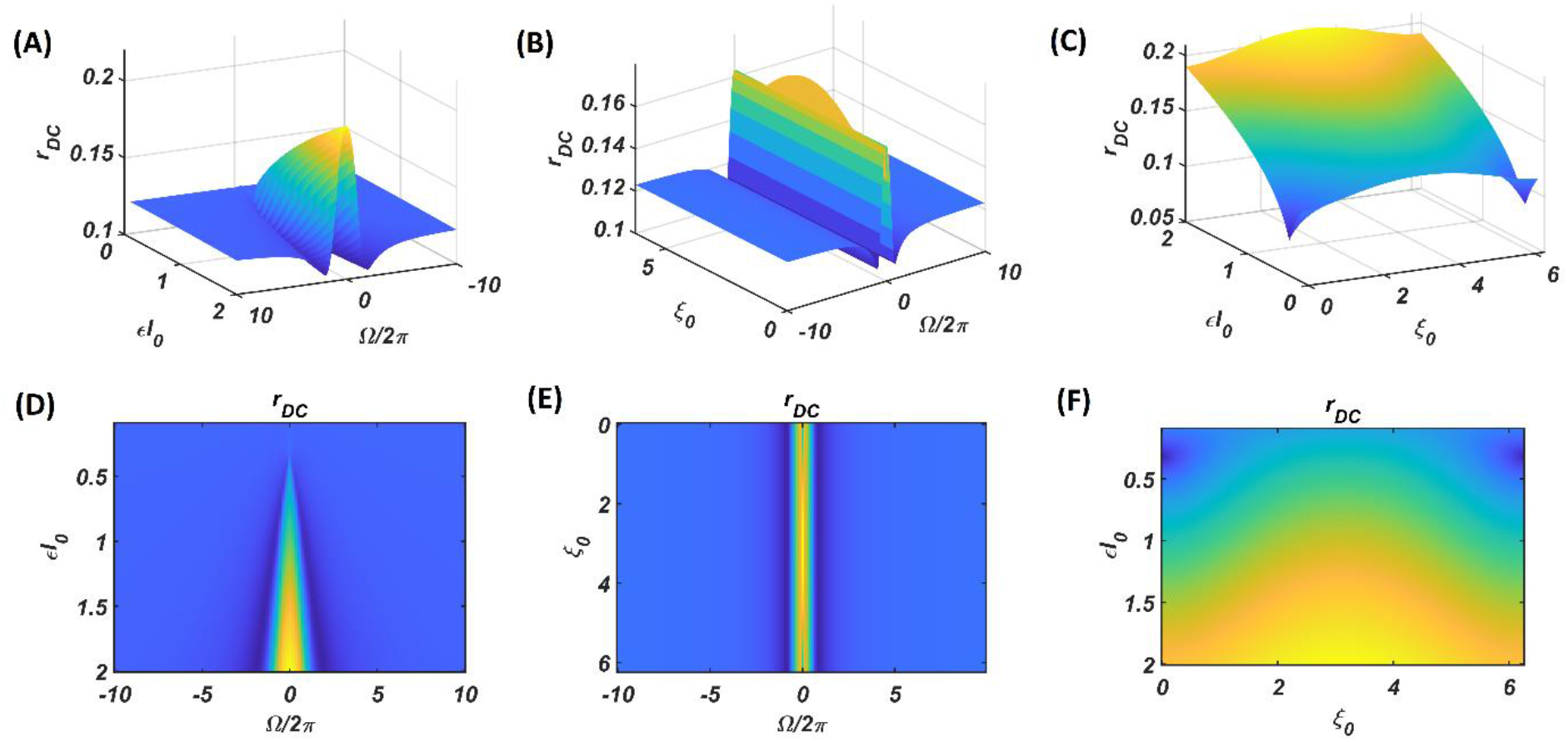
The DC response of the CAO oscillator w.r.t the three properties of the external input signal; amplitude (*εI*_0_), frequency (*ω*_0_) and phase offset (*ξ*_0_). For all these simulations the common parameters are: *μ* = 1, *β*_1_ = –150, *ω* = 2*π* × 60, *F* = 1, *μ_r_* = 1, *β*_1*r*_ = –10, *ω_r_* = 2*π*× 60, *A_r_* = 0.5, *θ_r_* = *π*. For the plots *ω*_0_ is varied from 2*π* × 50 to 2*π* × 70, *ξ*_0_ is varied from 0 to 2*π*, *ε* is varied from 0.1 to 2. For the plots in the left column *ξ*_0_ is kept fixed at a value of *π*. For the plots in the middle column *ε* is kept fixed at a value of 1. For the plots in the left column *ω*_0_ is kept fixed at a value of 2*π* × 60.

**Figure 13:**
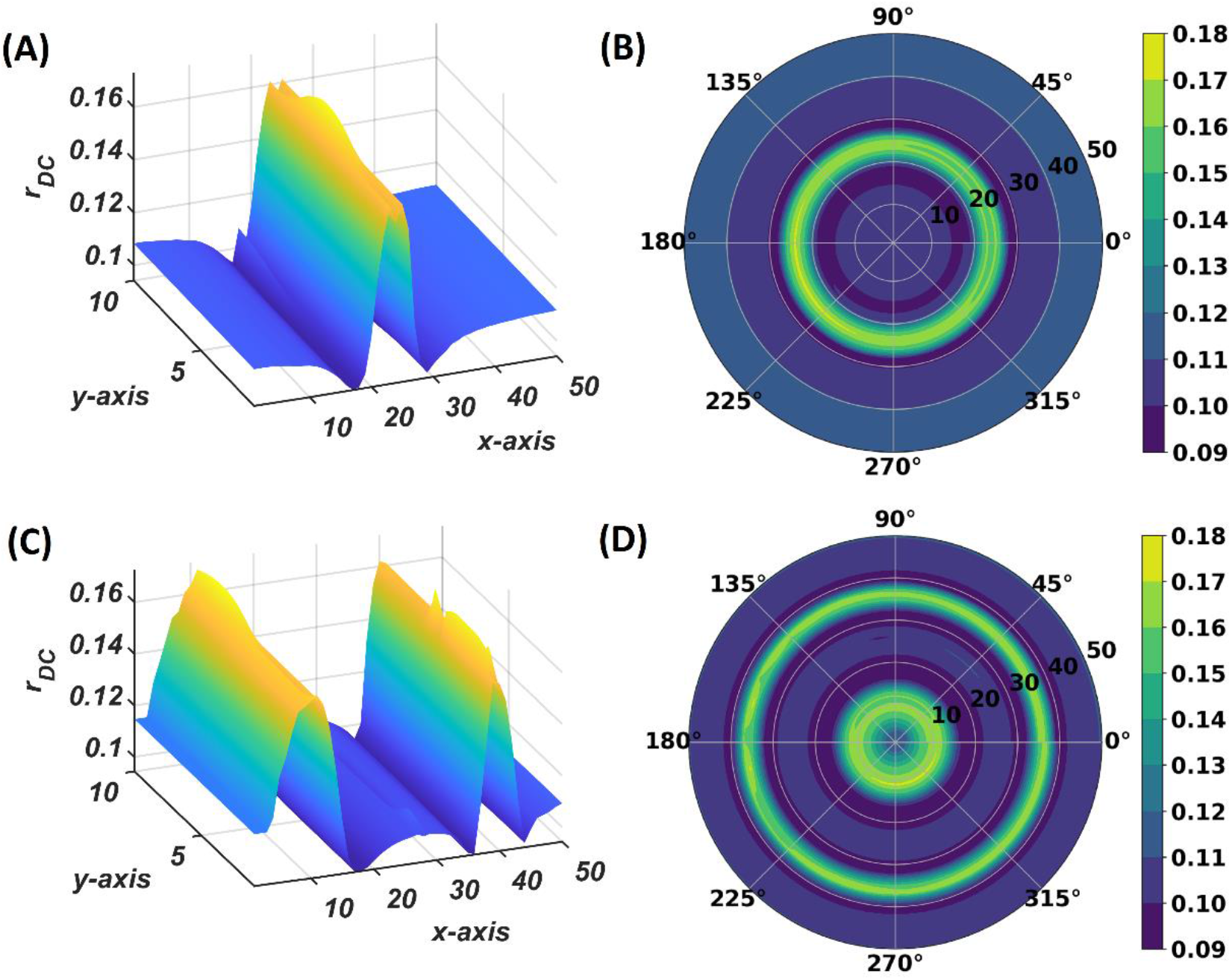
The subplots **(A)** and **(B)** depicts the response of the OTSOM model when the external input signal is: *I*(*t*) = cos(2*π* × 57.7029 + *π*). The subplots **(C)** and **(D)** depicts the response of the OTSOM model when the external input signal is: 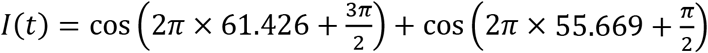. The *ω_ij_*’s of the CAO oscillators and the 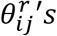 are adopted from the training stage of the OTSOM model as depicted in the fig. 10 and fig. 11C. The other parameters of the model are mostly preserved as used in the training session of the model: *μ* = 1, *β*_1_ = 150, *F* = 1, *μ_r_* = 1, *β*_1*r*_ = 1, *ω_r_* = 2*π* × 60, *A_r_* = 0.1.

The response of the OTSOM model is tested with three types of signals: periodic real sinusoidal signal (fig. 13A and 13B), quasi-periodic signal which is a combination two proximal frequency components (fig, 13C and 13D) given in the caption of fig. 13 and an aperiodic signal such as an Electroencephalograph (EEG) signal as plotted in fig. 14A with its characterizing power spectrum plotted in fig. 14B. The EEG signal was collected during mind wondering task with a sampling rate of 1024 Hz (Grandchamp et al., 2014). The characterizing *ω_ij_* and 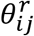 parameters during the testing phase is illustrated in fig. 10 and 12C respectively. It can be observed from fig. 13C and 13D that the representation of the frequency 55.669 Hz is broader w.r.t the frequency 61.426 Hz, which is because of the slope of *ω_ij_*s along the x-axis at around 55.669 Hz is lesser than the slope at around 61.426 Hz.

**Figure 14:**
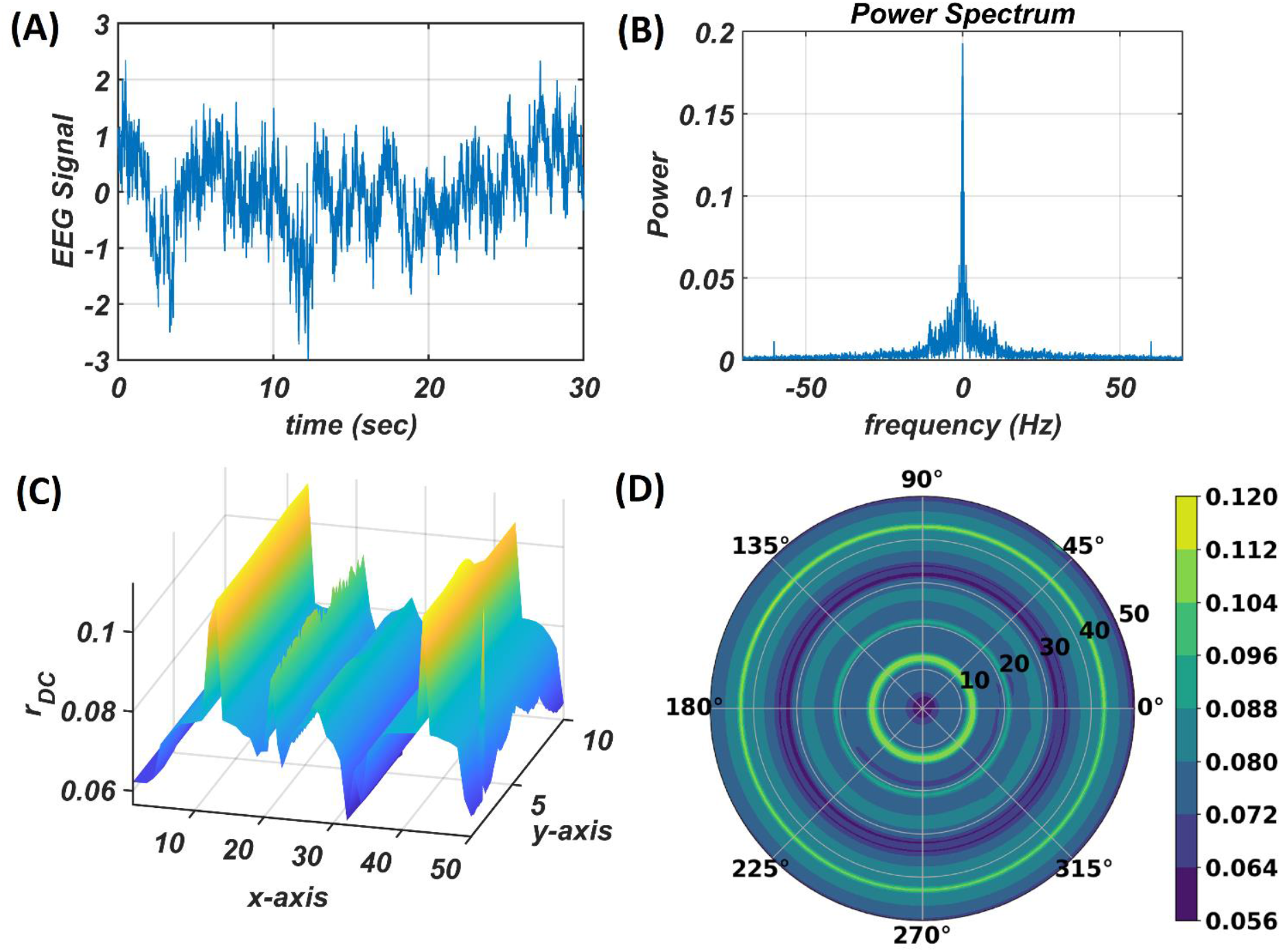
The subplots **(C)** and **(D)** illustrates the response of the OTSOM model when an arbitrary aperiodic signal such as the EEG signal plotted in **(A)** characterized by its power spectrum plotted in **(B)** is presented as external input signal.

## 4 Discussion

Tonotopic map refers to a map of tones or individual frequencies often found in auditory cortices of mammals. Optimal response at a specific frequency is characteristic of resonance. Based on this insight, we designed a tonotopic map model based on nonlinear oscillators capable of exhibiting resonance. We present a model of a tonotopic map, which consists of an array of Hopf oscillators, labeled as CAO. The map is trained on complex sinusoidal stimuli such that the frequency is mapped onto to the columns, and phase is mapped onto the rows. (The phase of the input signal is defined with reference to a reference oscillator labeled as SRO.) In other words, when a complex sinusoid with a given frequency and phase is presented as input stimulus, the oscillator at a specific row and column, whose frequency and phase are the closest to the input parameters, responds with the highest amplitude.

Existing computational models of tonotopic map do not attempt to model the underlying oscillation or the associated resonance in modeling tuned responses to pure tones. In the tonotopic model of (Ritter et al., 1992), which is based on a SOM model, frequencies are modeled as explicit parameters defined out of the context of the underlying oscillatory process. The model was able to achieve an ordered map of frequencies, with greater areas of the map differentially allotted to dominant frequencies in the input. However, the model was not able to capture any other temporal aspects of the input signal, since no signal was explicitly modeled. Another tonotopic map model described by Palakal et al., 1995 modeled the distribution of both frequency and time delay. But here too, these parameters are described as independent parameters, taken out of the context of the underlying temporal process. In this regard, the proposed tonotopic model based on oscillators and resonance represents a significant step forward.

A previous model (Biswas et al., 2021) that shows how a network of Hopf oscillators can be trained to learn arbitrary aperiodic signals was developed further to create the proposed tonotopic map model. To this end two improvements had to be made to the previous model:

a. a key element of (Biswas et al., 2021) is the concept of power coupling that achieves a stable (normalized) phase relationship between a pair of oscillators with arbitrary intrinsic frequencies. This scheme had to be modified in the proposed model since it must allow mixed forms of coupling combining power coupling with ordinary real coupling.
b. in the proposed model, oscillators must exhibit tuned responses not only to frequency but also to phase. In order to define a phase offset of the input signal, we introduced a reference oscillator (SRO) that projects to all the oscillators in the map.

The functional unit of OTSOM model is a single CAO oscillator that receives input from the external input and the SRO. We performed qualitative and the quantitative analysis of this unit under two conditions:

1. Magnitude of the SRO input is negligible compared to the external input.
2. Magnitude of the SRO input is comparable to the external input.

The magnitude of the input from the reference oscillator is 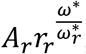, where the steady state value of the magnitude of oscillation of the SRO depends on *μ_r_* and *β*_1*r*_. As the SRO operates in the supercritical Hopf regime both *μ_r_* and *β*_1*r*_ are positive and the steady state magnitude of oscillation is 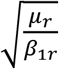. With this simple setup we have observed that the CAO oscillator can encode not only the frequency of the complex sinusoidal input signal primarily inside the entrainment regime, but also the phase offset of the input signal.

The entrainment regime of a canonical Hopf oscillator is previously analyzed by (Kim & Large, 2015) in terms of analyzing steady-state dynamical characteristics on *r* – *ψ* plane. As *ψ* is the angular difference between the oscillator and the external input signal, when the system exhibits stable fixed point (Δ> 0, *T*^2^ – 4Δ> 0, *T* < 0) or stable spiral behaviour (Δ> 0, *T*^2^ – 4Δ< 0, F*T* < 0) it can be interpreted that the system is entrained. When the relative phase of the oscillator w.r.t the input signal reaches a steady state value, it essentially means the actual frequency of oscillation of the oscillator is adapted from its natural frequency of oscillation to the frequency of the driving signal. Kim and Large (Kim & Large, 2015) have analyzed the effect of the strength of the driving signal on its entrainment characteristics by mapping the nature of the steady-state solution on the *εF* vs Ω space. There are five possible steady state solutions exhibited by four regimes of canonical Hopf oscillator defined by its intrinsic parameter values. These five steady state solutions are stable node, stable spiral, unstable node, unstable spiral and saddle point. When the Hopf oscillator operates in critical Hopf parameter regime (*μ* = 0, *β*_1_ > 0, *β*_2_ = 0) it exhibits either stable node or stable spiral solution at steady-state, i.e., for any values of its intrinsic parameter *β*_1_, the strength of the driving signal (*εF*) and Ω it is going to be entrained to the frequency of the driving signal. Therefore, it can be stated that the entrainment regime of the Hopf oscillator operating in the critical parameter regime is unbounded. In both of these cases the state of the system reaches the fixed point asymptotically, i.e., it takes forever for the oscillator to get entrained. Generally, a small neighbourhood around the fixed point is defined to declare the entrainment of the system. The Hopf oscillator operating in the supercritical parameter regime (*μ* > 0, *β*_1_ > 0, *β*_2_ = 0) exhibits three steady-state solutions: stable fixed point, stable spiral and unstable spiral. Till the boundary to the unstable spiral solution the system exhibits entrainment.

The typical initial value of the variance *η_ω_* has the property: *σ_yωm_* ≫ *σ_xωm_*. Due to high variance along the column, the other oscillators in the same column as the winner tend to adapt to the feature of the presented input pattern at the same rate as the winner neuron, which ensures the low variability of the learnt natural frequency of the oscillator along y-axis. An initial standard deviation of *σ_yωm_* = 100 and *σ_xωm_* = 4 is sufficient for the tonotopic organization to arise as presented in fig. 10. Although there is a lower bound for *σ_yωm_* depending on the number of oscillators along the *y*- direction, *N_y_*, there are no strict bound on *σ_xωm_* depending on the dimensionality of the 2D array of oscillators. It can be interpreted that the lower bound on *σ_yωm_* should be proportional to *N_y_* as the greater the number of oscillators per column, the lesser the adaptability rate of the oscillators at the boundaries of the adaptable neighbourhood along the column. However, a square neighbourhood window function is chosen for the simulation presented in this study, a rectangular window function is also feasible. A rectangular window function can be defined by *d_r_* ≠ *d_c_*. *σ_xω_* and *σ_yω_* decrease in a Gaussian contour w.r.t time at a much slower time scale to model the effect of annealing. *σ_xω_* and *σ_yω_* are updated after every epoch, with a typical standard deviation on an iterative time scale of 500*n_epft,ω_*.

A few aspects need to be elaborated about the 2^nd^ stage of training. The 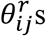 were failing to self-organizing themselves in a linearly increasing or decreasing fashion along the column when the CAO oscillators were placed on a 2-dimensional rectangular grid. To fix this issue, periodic boundary condition is introduced along the spatial dimension of the column i.e., the bottom row of the CAO is closest with an equidistant to both the top row as well as the second last row which is the motivation behind representing the CAO on a polar coordinate representation. Although this ensured that the 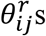 self-organize themselves in a linearly increasing or decreasing manner in a given column, 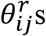 were slipping at a constant rate along the azimuth axis which can be observed in fig. 15A. To fix this problem the 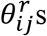 of the top most row of the CAO were fixed at 0° angle and from fig. 15B it can be observed that 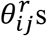 of a given column were stabilized with a linear organization. On the contrary, the *ω_ij_*s are able to self-organize themselves without the aforementioned periodic boundary condition.

**Fig15:**
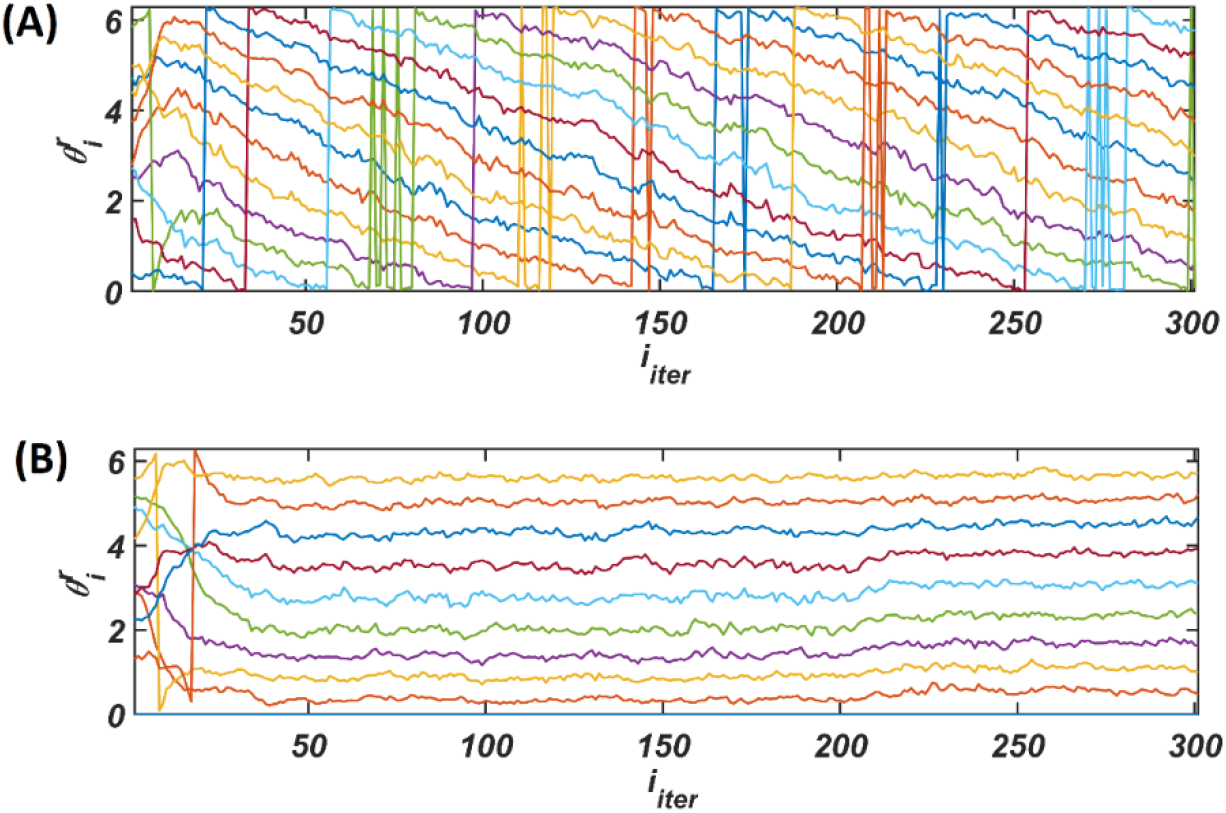
**(A)** Evolution of the 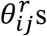 of a given column over multiple epochs in the 2^nd^ stage of training with the periodic CAO along the dimension of y-axis, **(B)** Self organization of 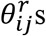 of the same column throughout the 2^nd^ stage of training with the periodic CAO along the dimension of y-axis along with fixed 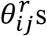 of the top most row with 0° angle.

A Comparison with the conventional SOM: For conventional SOM model (Kohonen, 1998), the neuronal response is characterized by its linear or nonlinear activation function. When these rate coded neurons are a part of SOM framework, the afferent weights for a particular neuron are also considered to be an internal feature of the neurons, considering close proximity of these afferent synapses to the corresponding neurons. The key differences between the conventional SOM model and OTSOM are:

1. the afferent connection weights are fixed,
2. the input is time varying complex sinusoidal signal instead of a constant vector,
3. the neurons are limit cycle oscillators instead of rate-coded neurons with a static transfer function.

Although we have tested the model with complex sinusoidal input signals sampled from the frequency band from 55 to 65 Hz, the bandwidth can be scaled up/down or shifted. The proposed model can be used to explain the tonotopic organization evolved in auditory cortex of mammals.

## 6 Author Contributions

DB and AT: hypothesis testing, conceptualization, theory development, numerical simulations, investigation, methodology, and validation. DB and VC: visualization and writing—original draft. VC: writing—review, editing, and supervision. All authors contributed to the article and approved the submitted version.

## 7 Data availability statement

The original contributions presented in the study are included in the article/Supplementary Material, further inquiries can be directed to the corresponding author/s.

## 9 Acknowledgments

We acknowledge the support of fellow lab mate Sayan Ghosh for preprocessing the EEG signals

## Appendix 1: A pair of Hopf oscillators coupled bilaterally and unilaterally through modifier power coupling

The dynamics of a pair of Hopf oscillators bilaterally coupled through modified power coupling is described by the eqns. 6, 7 and 8. Whereas the dynamics of a pair of Hopf oscillators unilaterally coupled through modified power coupling which is identical as the single unit of the OTSOM model under the special condition *ε* ≅ 0 is described by eqns. 12, 13 and 14. The schematic of these networks are shown in the fig. A1.

**Figure A1:**
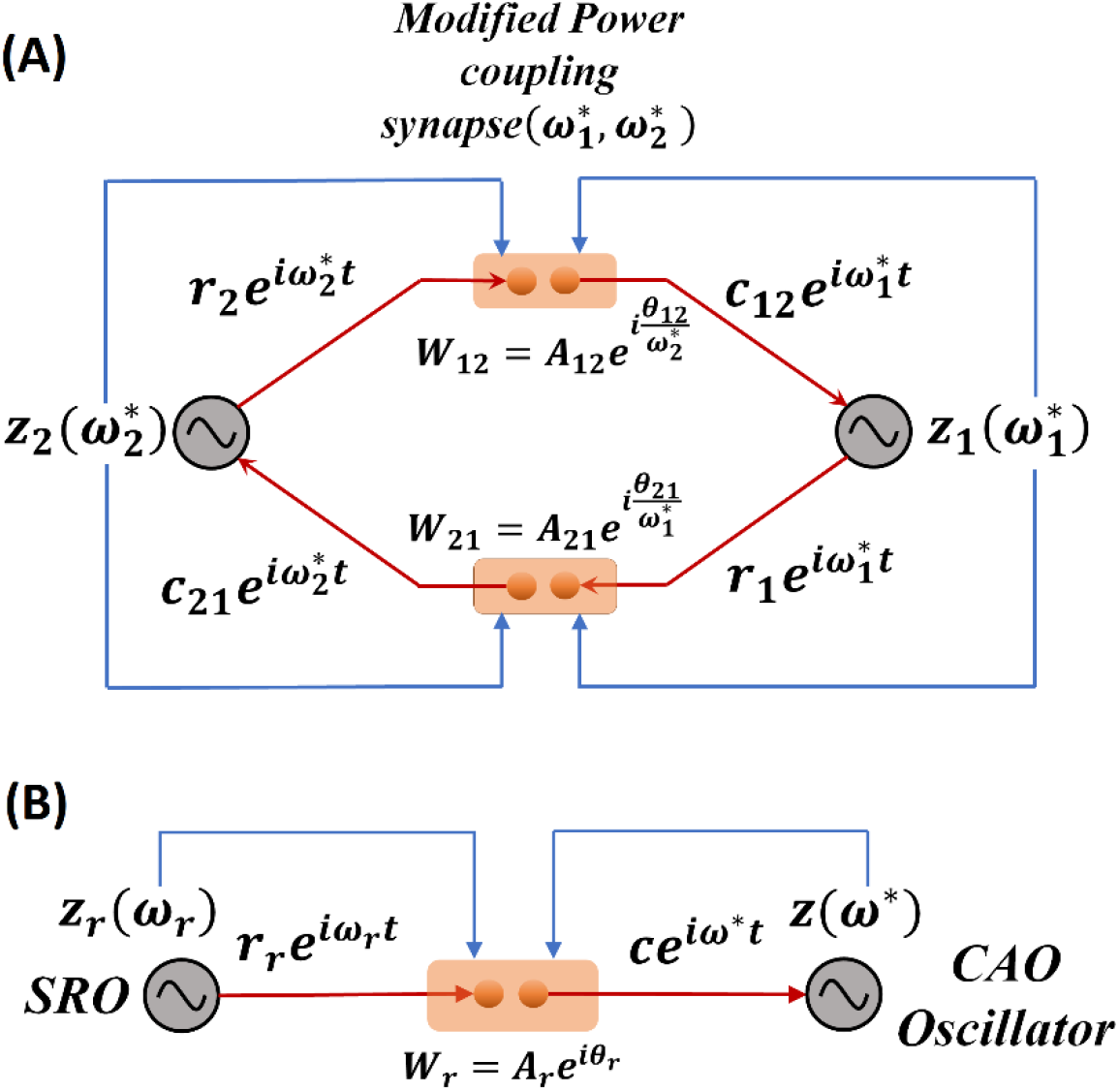
Network schematic of a pair of Hopf oscillators bilaterally **(A)** and unilaterally **(B)** coupled through modified power coupling.

### Network 1 (bilateral coupling)

Initially both the oscillators will receive complex sinusoidal input signal from the other oscillator through modified power coupling connection as 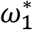 and 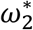 are initialised to *ω*_1_ and *ω*_2_ respectively. As the Hopf oscillator keeps oscillating at its natural frequency, *ω*, when it is perturbed by a complex sinusoidal signal with the same frequency, *ω*_0_ = *ω*, both of the oscillators in the pair will continue to oscillate at a same frequency. So, 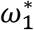 and 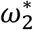 will remain as *ω*_1_ and *ω*_2_. Defining the new normalized phase difference as, 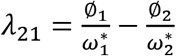, assuming at steady state 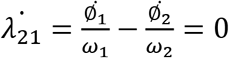, considering 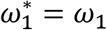 and 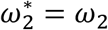 all along. From equation 2b and 3b,

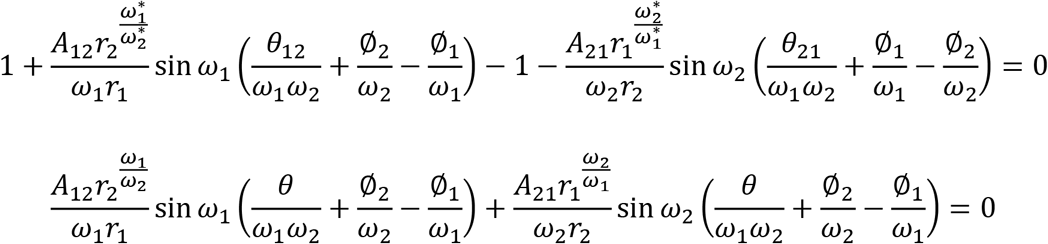

As we have already assumed, *θ*_12_ = –*θ*_21_ = *θ*, it is obvious that for some of the solutions of the above equation both the terms in the L.H.S of the equation would be 0, in that case;

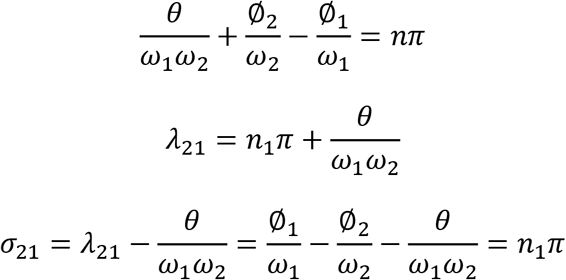

where, *n*_1_ = –*n*. So. The normalized phase difference has multiple solutions which depends on *θ* as well as the natural frequency of both of the oscillators.

### Network 2 (unilateral coupling)

Initially the CAO oscillators will receive complex sinusoidal input signal from the SRO through modified power coupling connection as *ω*\* is initialised to *ω*. As the Hopf oscillator keeps oscillating at its natural frequency, *ω*, when it is perturbed by a complex sinusoidal signal with the same frequency, *ω*_0_ = *ω*, CAO oscillator will continue to oscillate at a same frequency. So, *ω*\* will remain as *ω*. The normalized phase difference between CAO oscillator and SRO will be:

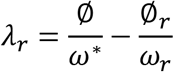

As *ω*\* will remain *ω* the normalised phase difference will be: 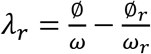. Assuming 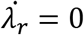 at steady-state;

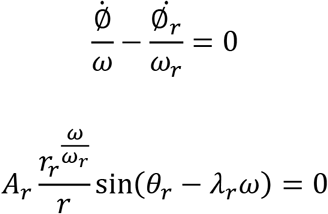

The only solution of which is: 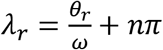. So, the pair of unilaterally coupled Hopf oscillators through modified power coupling will always synchronize with each other with a normalized phase difference of 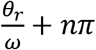.

## Appendix 2: The steady state dynamical analysis of single unit as the CAO oscillator operates under the entrainment regime of the input signal

The approximate dynamics of eqns. 12, 13, 14 considering *A_r_* ≪ *εI*_0_, when *ω* = *ω*_0_:

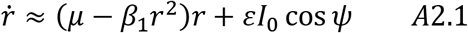

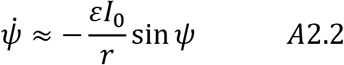

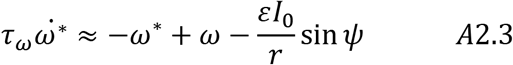

At steady state, 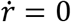, 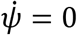. From equation A2.2:

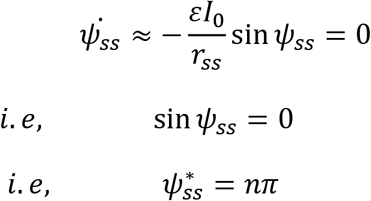

The solutions 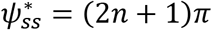 are unstable whereas the solutions 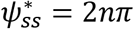 are stable. For the stable solutions;

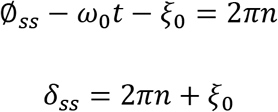

i.e., the steady state phase offset of the CAO oscillator will be same as the phase offset of the input signal. For 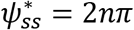 solutions, equation A2.1 will become:

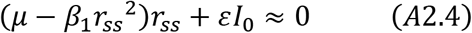

From the last expression, the positive real solution, *r_ss_*, of (A2.4) is derived with the help of MathWorks equation solver:

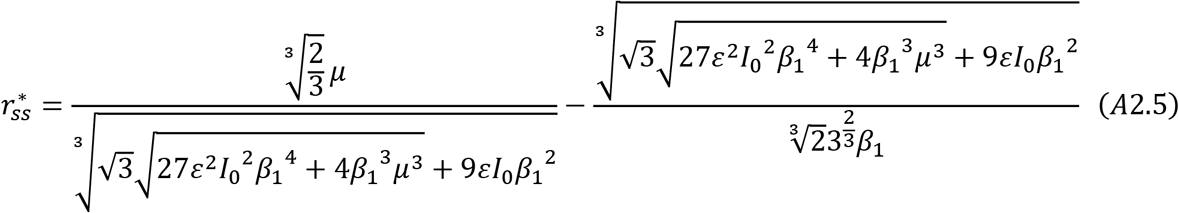

The solution of steady state magnitude of oscillation as found in eqn. A2.5 is numerically verified in fig. A2.1. Whereas the fig. A2.2 elaborates the dependency of the entrainment width and the typical transient time on the parameters *μ, β*_1_ and *εI*_0_.

**Figure A2.1:**
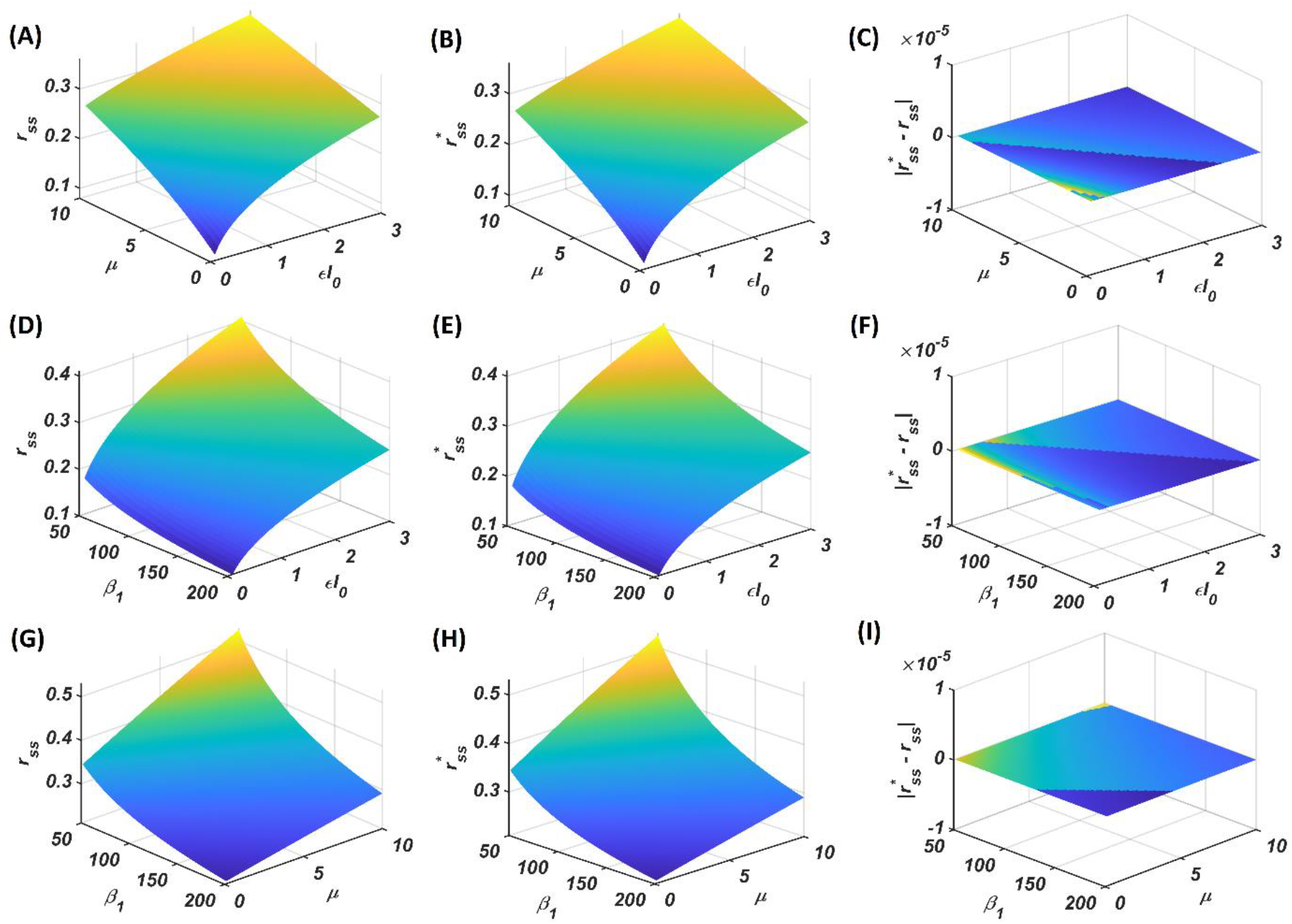
The first two columns (subplots **A, D, G** and **B, E, H**) proclaim that the dependency of the solution 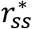 of the polynomial expression given by eqn.-A2.4 found by MathWorks polynomial solver, given by the expression in eqn.-A2.5 on two of the three parameters *μ, β*_1_ and *εI*_0_ keeping the remaining fixed at a certain intermediate value (*εI*_0_ = 2, *ξ*_1_ ² 150, *μ* = 1) is identical to the steady state magnitude of oscillation (*r_ss_*) deduced by simulating the transient dynamics of eqns.- A2.1 and A2.2 with *ω* = *ω*_0_ = 2*π* × 60. The error between the first two columns is plotted in the 3^rd^ column (subplots **C, F, I**).

**Figure A2.2:**
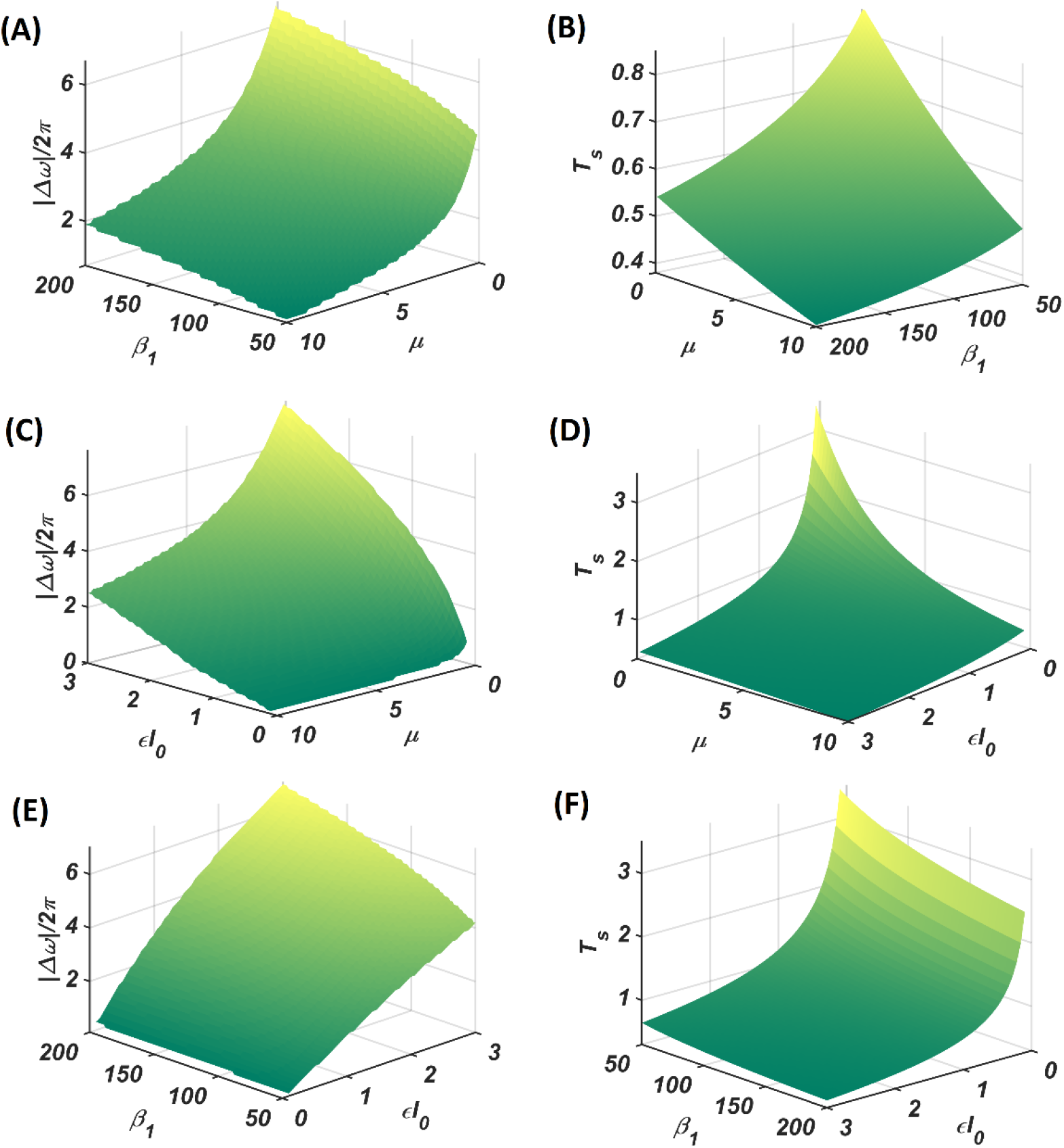
The subplots **A**, **C**, **E** delineates the dependency of the width of the entrainment regime (Δ*ω*) on two of the three parameters *μ, β*_1_ and *ε* keeping the remaining fixed at a certain intermediate value, similarly the remaining subplots delineates the dependency of *T_s_* (the time an individual oscillator takes to attain steady-state under the influence of complex sinusoidal external input signal). Parameter value: For **(A)** and **(B)** *εI*_0_ = 2, for **(C)** and **(D)** *β*_1_ = 150 and for **(E)** and **(F)** *μ* = 1, *ω* is kept fixed at 2*π* × 60 rad/sec while *ω*_0_ is varied from 2*π* × 45 rad/sec to 2*π* × 75 rad/sec.

The approximate dynamics when the CAO oscillator is under the entrainment regime with *ω* ≠ *ω*_0_:

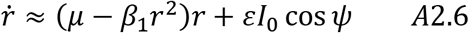

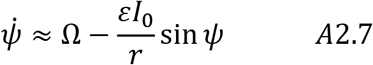

At steady state, equating eqn. A2.7 to zero;

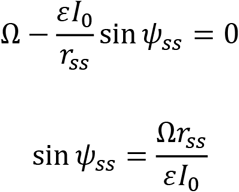

Substituting cos *ψ_ss_* into eqn. A2.4 with the steady state assumption,

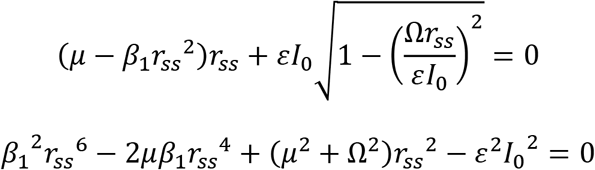

The solution of the expression can be drawn using MathWorks polynomial solver. The only positive real solution of the equation is:

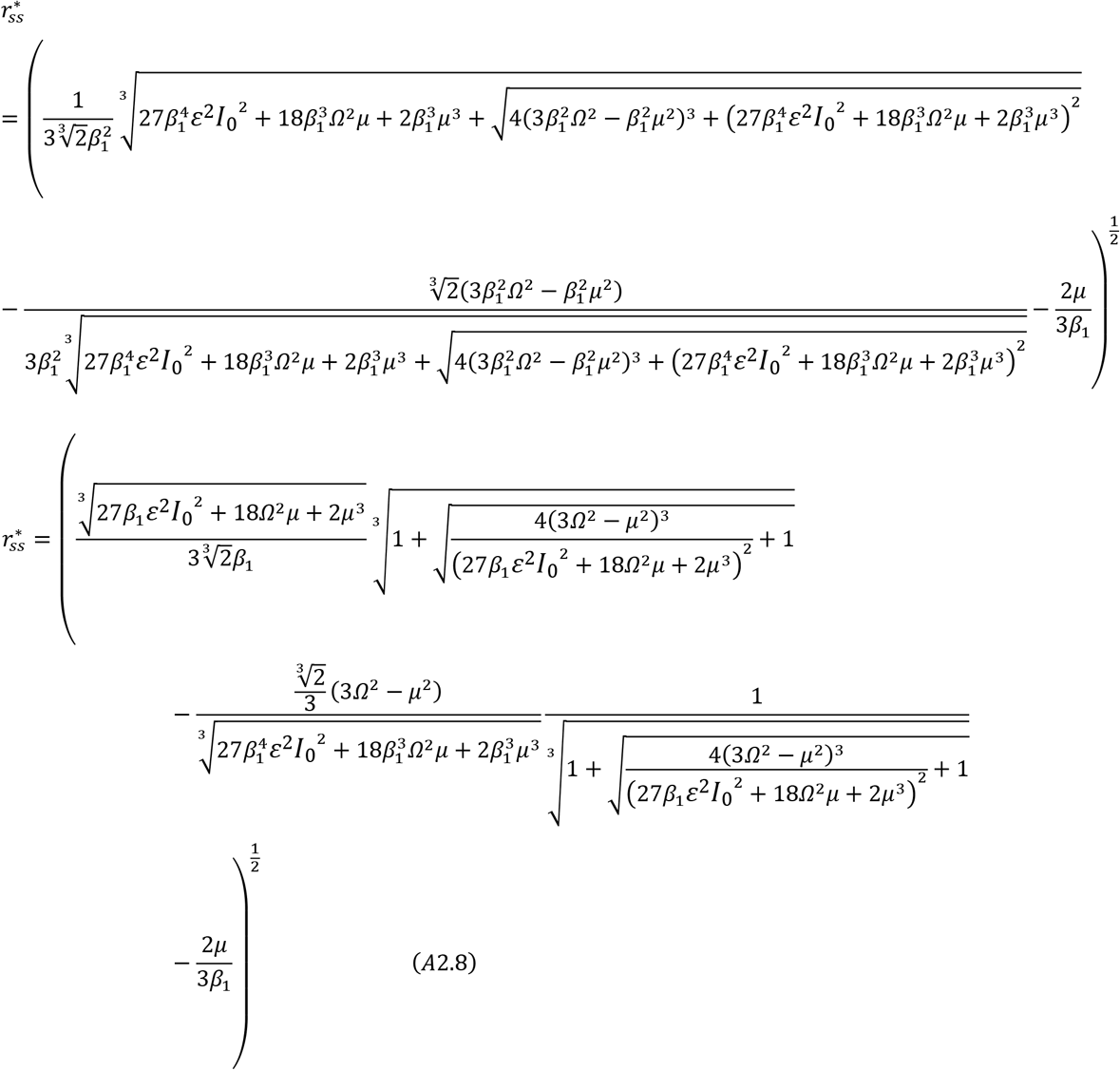

Assuming entrainment, the steady state phase offset of the main oscillator can be derived as follows:

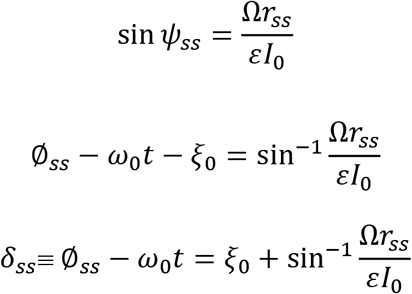

## Appendix 3: Steady state dynamics of adaptive Hopf phase of first stage of training

The dynamics of a single CAO oscillator, in polar coordinates, can be approximated for the following condition (*A_r_* ≪ *εI*_0_):

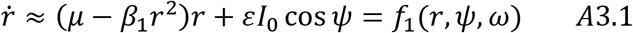

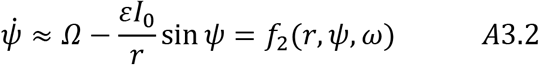

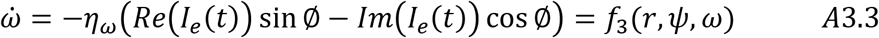

where, *I_e_*(*t*) = *εI*(*t*) = *εI*_0_*e*^*i*(*ω*_0_*t*+*ξ*_0_)^, the *ω* dynamics can be simplified as following:

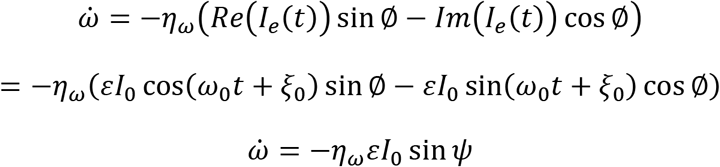

At steady state, 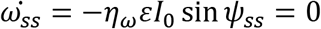. The stable solution of which would be, 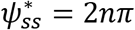. Similarly, from eqn. A3.2,

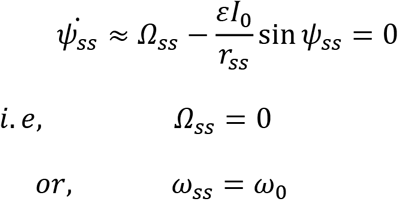

So, the steady state magnitude of oscillation will be the solution of the following expression as described in Appendix-2:

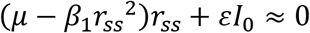

In this scenario the phase offset of the CAO oscillator at steady state will be:

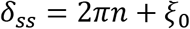

The corresponding Jacobian matrix for eqns. (A3.1-A3.3):

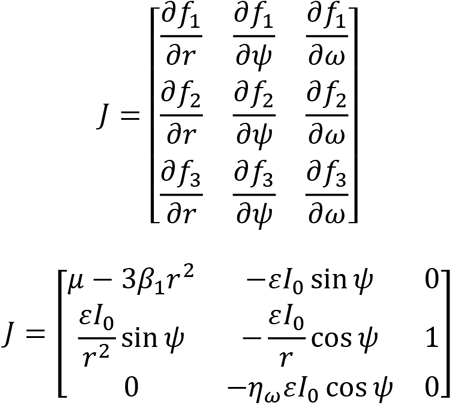

The Jacobian matrix at the fixed point or the steady state solution:

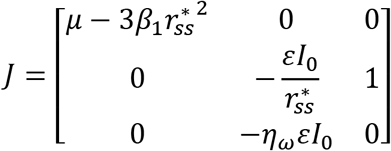

Assuming *α*_1_ < 0, the determinant (Δ) and the trace (*T*) respectively:

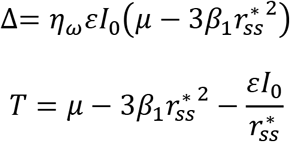

So, the solution 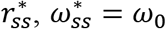 and 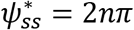 will be saddle point as 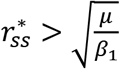, which ensures 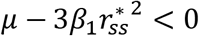, or Δ< 0.

## Appendix-4: Steady state dynamics of the transient phase of the second stage of training

The dynamics of the single unit of the network during the transient phase of the 2^nd^ stage of training is defined by eqns. 12, 13, 14. The complex variable counterpart of eqns. 12 and 13 are:

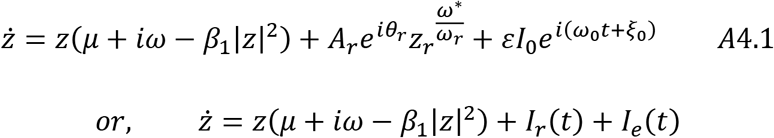

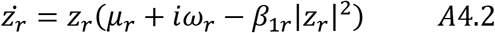

The complex activation of the reference oscillator at steady state,

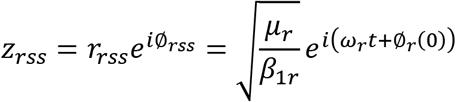

Assuming the special condition, *Ω* < Δ*ω*, *ω* ≠ *ω*_0_ and *ϵ_max_* > *εI*_0_ – > *ϵ_min_* i.e., the CAO oscillator is operating under the entrainment regime of the input signal (where Δ*ω* depends on the *μ, β*_1_, *ε, F, ξ*_0_, *θ_r_* and *A_r_*) with a visible amount of interference between *I_e_*(*t*) and *I_r_*(*t*), at steady state, the frequency of the CAO oscillator becomes the frequency of the input signal i.e., 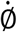 or *ω*\* becomes equal to *ω*_0_. So, the steady state version of eqn. A4.1 can be written as:

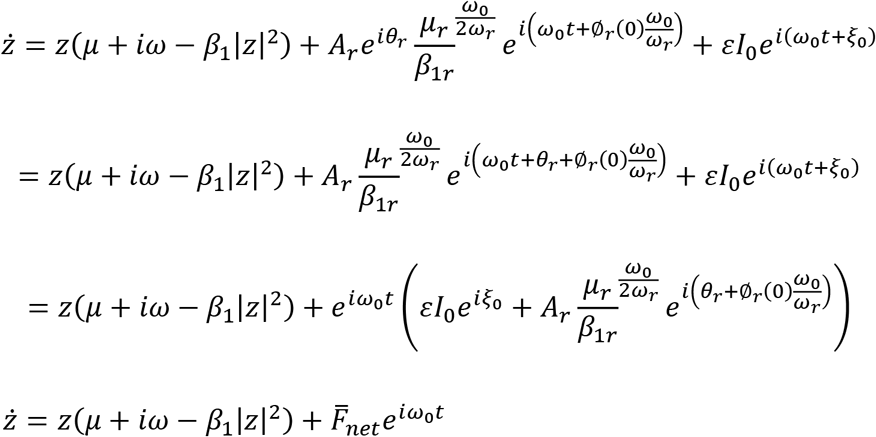

where,

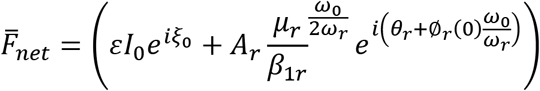

is the phasor corresponding to the net input to the CAO oscillator at steady-state. The magnitude and the phase offset of 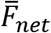 are:

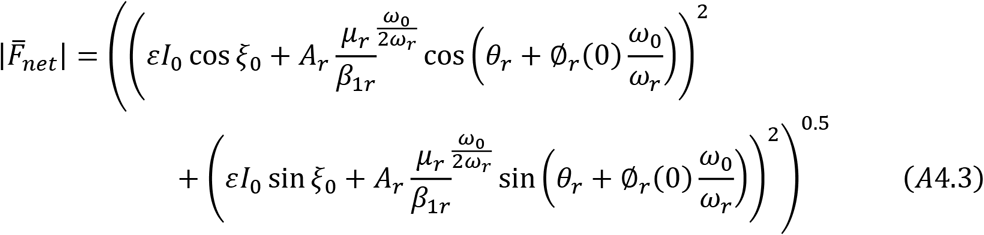

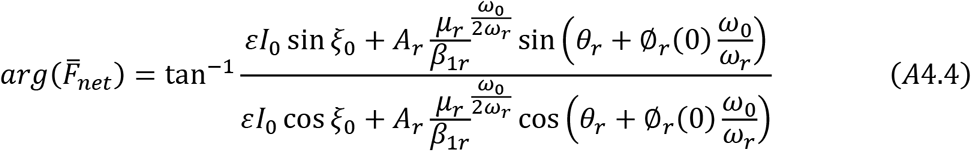

The steady-state magnitude of oscillation is dependent on 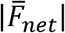 whereas the phase offset of oscillation 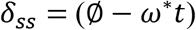 will be same as 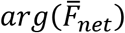 which is verified in fig. 10C.

## Appendix-5: Steady state dynamics of Hebbian plasticity phase of second stage of training

The approximate dynamics of the single unit at Hebbian plasticity phase of the 2^nd^ stage with the assumption *εI*_0_ ≫ *A_r_*:

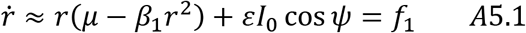

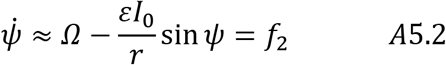

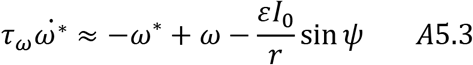

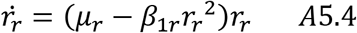

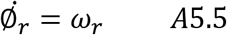

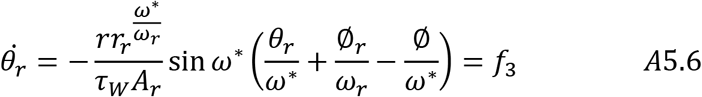

Considering the CAO oscillator is operating inside the entrainment regime, at steady state *ω*\* becomes *ω*_0_. From equation-A5.2 we get;

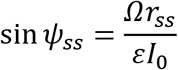

So, 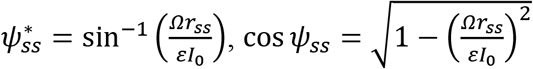 and the phase offset of the CAO oscillator at steady-state 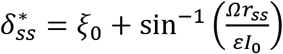. The steady state dynamics of equation A5.6:

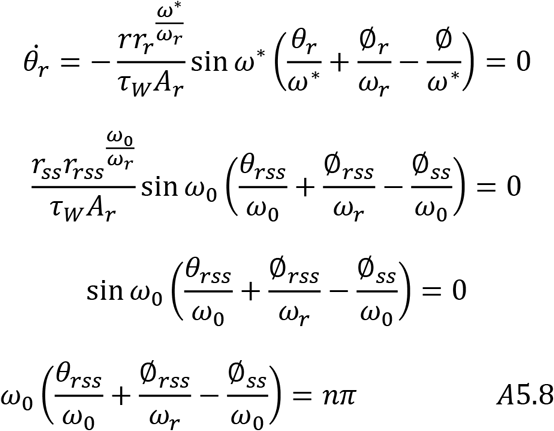

At steady state, 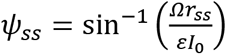 or, 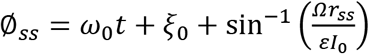, and 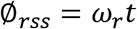, assuming 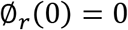, substituting the values of 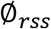 and 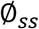 into equation A4.8.

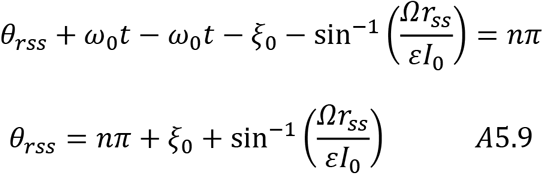

Likewise, 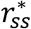 would be the solution of the following expression:

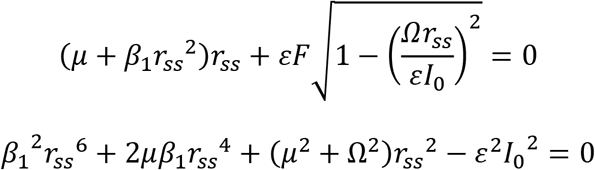

## Appendix-6: Steady state dynamics of Hebbian plasticity phase of 2^nd^ stage of training under the condition *A_r_* ≠ 0

Polar coordinate representation of the dynamics:

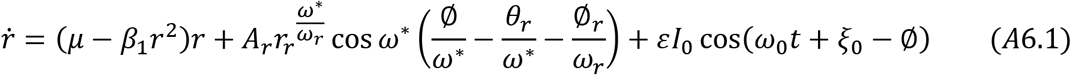

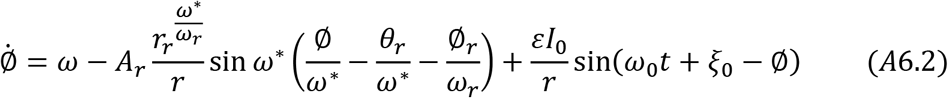

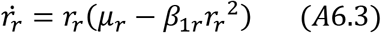

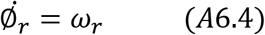

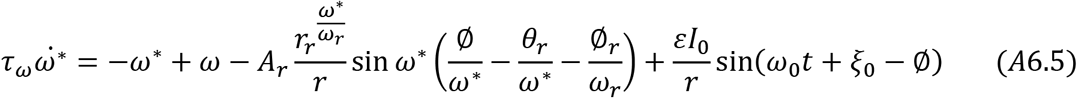

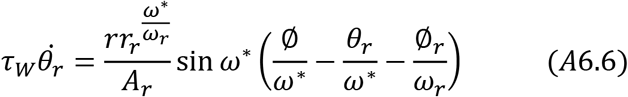

A brief analysis of the steady state dynamics as follows:

Considering the CAO oscillator is operating inside the entrainment regime, at steady state *ω*\* becomes *ω*_0_. Writing eqns. A6.1 and A6.2 in terms of *ψ* and *Ω*.

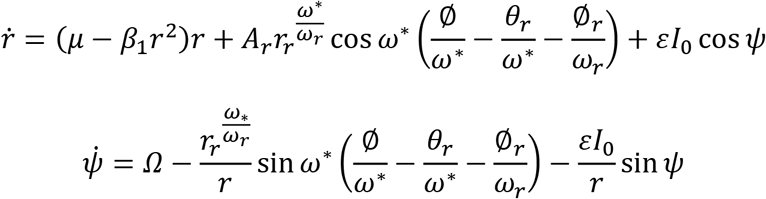

In the entrainment regime, *ω*\* becomes *ω*_0_ at steady state.

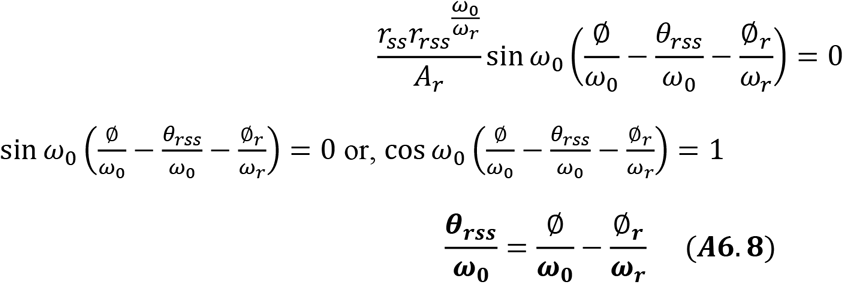

From eqn. A6.6 at steady state:

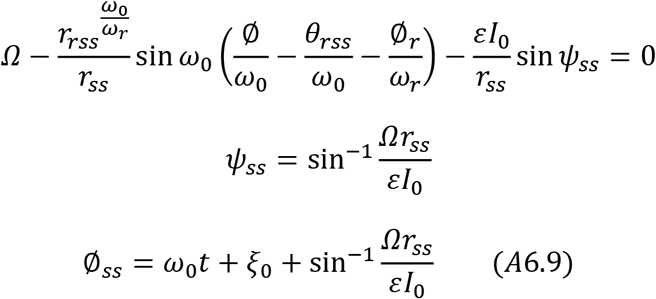

From eqns. A6.8 and A6.9 assuming the reference oscillator was initialized at 0 phase, 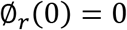.

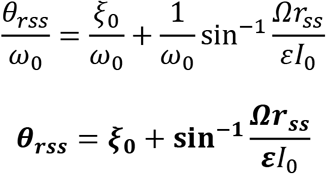

From eqn. A6.1 we can find the steady state value of *r* by finding the solution of the following equation;

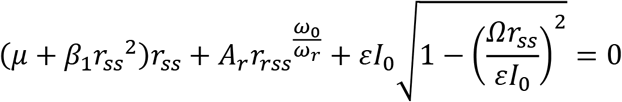

